# Small interfering RNAs based on huntingtin trinucleotide repeats are highly toxic to cancer cells

**DOI:** 10.1101/247429

**Authors:** Andrea E. Murmann, Quan Q. Gao, William Putzbach, Monal Patel, Elizabeth T. Bartom, Calvin Law, Bryan Bridgeman, Siquan Chen, Kaylin M. McMahon, C. Shad Thaxton, Marcus E. Peter

## Abstract

Trinucleotide repeat (TNR) expansions in the genome cause a number of degenerative diseases. A prominent TNR expansion involves the triplet CAG in the huntingtin (HTT) gene responsible for Huntington’s disease (HD). Pathology is caused by protein and RNA generated from the TNR regions including small siRNA-sized repeat fragments. An inverse correlation between the length of the repeats in HTT and cancer incidence has been reported for HD patients. We now show that siRNAs based on the CAG TNR are toxic to cancer cells by targeting genes that contain long reverse complimentary TNRs in their open reading frames. Of the 60 siRNAs based on the different TNRs, the 6 members in the CAG/CUG family of related TNRs are the most toxic to both human and mouse cancer cells. siCAG/CUG TNR-based siRNAs induce cell death *in vitro* in all tested cancer cell lines and slow down tumor growth in a preclinical mouse model of ovarian cancer with no signs of toxicity to the mice. We propose to explore TNR-based siRNAs as a novel form of anti-cancer reagents.

---

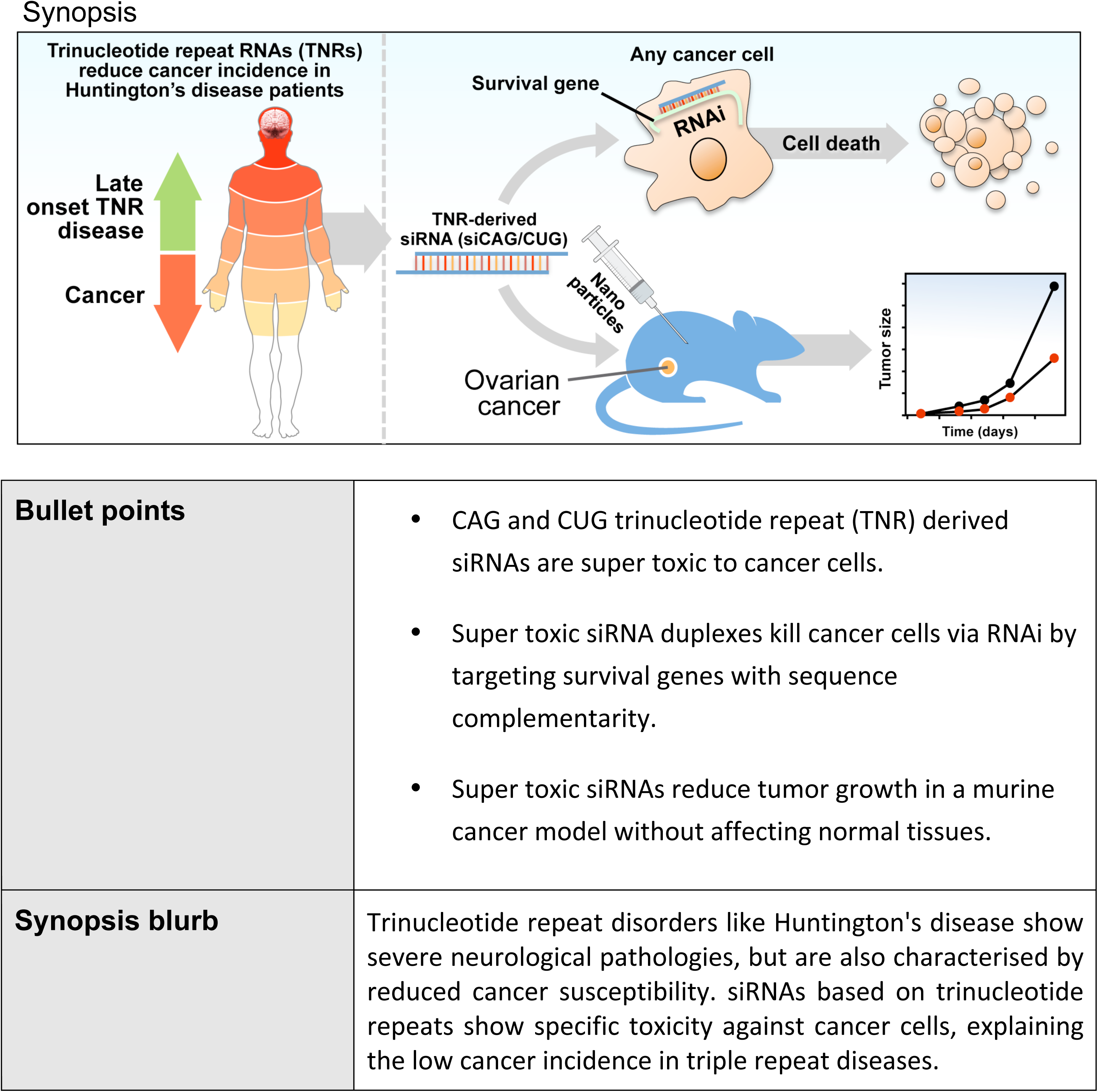

## Introduction

Trinucleotide repeat (TNR) expansions are the cause of a large number of degenerative disease syndromes characterized by amplification of DNA triplet motifs [1]. They include spinocerebellar ataxias (SCAs), spinobulbar muscular atrophy (SBMA), myotonic dystrophy type 1 (DM1), and Huntington’s disease (HD) [1, 2]. HD is a dominantly inherited neurodegenerative disorder caused by expansion of CAG repeats in the huntingtin (HTT) gene. It has been shown that the resulting glutamine expansions (polyQ) in HTT are toxic to cells [3, 4] and that the length of the CAG amplifications determines severity and onset of the disease [2, 4]. In addition to polyQ toxicity, repeat-associated, non-ATG translation (RAN translation) was discovered as another translation-level pathogenic mechanism of CAG repeat-containing mRNAs [5]. More recent evidence however, also points toward RNA playing a role in affecting cell viability by poly-triplet repeats [1, 6]. Indeed, many of the repeats in several TNR diseases are not located in open reading frames (ORFs) but in introns or untranslated regions (UTRs) [4]. DM1 is the best-characterized disease regarding RNA toxicity. The CUG repeats are in the 3’UTR of the dystrophia myotonica protein kinase (DMPK) gene, causing most of their toxicity by forming hairpin structures [7]. These hairpins are believed to recruit a number of RNA-binding proteins to nuclear RNA foci [8]. Another mechanism by which CAG/CUG TNRs could be toxic at the RNA level is by interfering with cellular splicing. This has been shown for CUG in DM1 [9] and CAG in HD [10].

Mounting evidence suggests the CAG TNR expansions are toxic at the RNA level. It was shown in *Drosophila* that the toxicity of the CAG repeat disease gene spinocerebellar ataxia type 3 (SCA3) protein ataxin-3, is in large part caused by the trinucleotide repeat RNA and not the polyQ protein [11]. Replacing some of the glutamine coding CAG repeats with the other codon coding for glutamine, CAA, mitigated the toxicity despite similar polyQ protein expression levels. Direct toxicity of mRNA with extended CAG repeats was also demonstrated in mice [12]. Finally, there is convincing evidence that CAG/CUG repeats can give rise to RNAi-active small RNAs. In human neuronal cells, expression of the CAG expanded exon 1 of HTT (above the threshold for complete penetrance which is > 40) [6] caused an increase in small CAG repeat-derived RNAs (sCAG) of about 21 nt in length. Above a certain length, CAG/CUG repeats were found to be cleaved by Dicer,the enzyme that generates mature miRNAs from pre-miRNAs before they are incorporated into the RNA induced silencing complex (RISC) [13]. The CAG repeat derived fragments could bind to complementary transcripts and downregulate their expression via an RNAi-based mechanism. In a mouse model of HD treatment of the mice with a locked nucleic acid–modified 20mer antisense oligonucleotide complementary to the CAG TNR (LNA-CTG) which reduced the expression of sCAGs but not of HTT mRNA or protein reversed motor deficits [14]. This study identified sCAG as a disease causing agent. Since sCAGs, isolated from HD human brains, when transfected reduced viability of neurons [6], these sequences might affect cell viability through RNAi by targeting genes that regulate cell survival.

We recently reported that si- and shRNAs derived from CD95, CD95L [15], and other genes in the human genome [16] kill cancer cells through RNAi by targeting a network of critical survival genes [15]. DISE (death induced by survival gene elimination) was found to involve simultaneous activation of multiple cell death pathways, and cancer cells have a hard time developing resistance to this form of cell death [17]. DISE was found to preferentially affect transformed cells [17]. Because the length of the CAG repeats in different CAG repeat diseases has been inversely correlated with cancer incidence in various organs [18–21], we were wondering whether RNAi active CAG based TNRs might be responsible for this phenomenon and whether they could be used to kill cancer cells.

We have now identified an entire family of TNR-based siRNAs - which contains the CAG repeat that causes HD - to be at least 10 times more toxic to cancer cells than any tested DISE-inducing si/shRNA. Our data suggest this super toxicity is caused by targeting multiple complementary TNR expansions present in the open reading frames (ORFs) of multiple genes, rather than in their 3’UTRs. As a proof of concept, we demonstrate that siCAG/CUG can be safely administered to mice to slow down growth of xenografted ovarian cancer cells with no obvious toxicity to the animals. We are proposing to develop super toxic TNR expansion-based siRNAs for cancer treatment.

## Results

### siCAG/CUG kills all cancer cells *in vitro*

CAG repeats are the defining factor in Huntington’s disease, and their complement CTG is amplified in myotonic dystrophy type 1 (DM1) [1]. We were interested in determining whether a 19mer duplex of CAG and CUG repeats (siCAG/CUG) (Fig 1A) would affect the growth of cancer cells. When transfecting siCAG/CUG into various human (Fig 1B) and mouse (Fig 1C) cancer cell lines at 10 nM, all cancer cells stopped growing within hours of transfection and eventually most of the cells died with no outgrowth of recovering cells (**Movies EV1-EV10**) (Appendix Fig S1A). All cancer cells transfected with siCAG/CUG showed morphological changes similar to the ones we observed in cells undergoing DISE (Appendix Fig S1B, [15, 17]). We found that siCAG/CUG killed HCT116 cells even when transfected at 10 pM (Fig 1D). Compared to any other si- or shRNA we have tested siCAG/CUG is ~10-100 times more toxic depending on the assay used. When monitoring cell viability (ATP content), the IC50 for siL3, the most toxic DISE inducing siRNA we have used, was determined to be 0.8 nM and for siCAG/CUG was 0.039 nM (Fig 1E).

**Figure 1.**
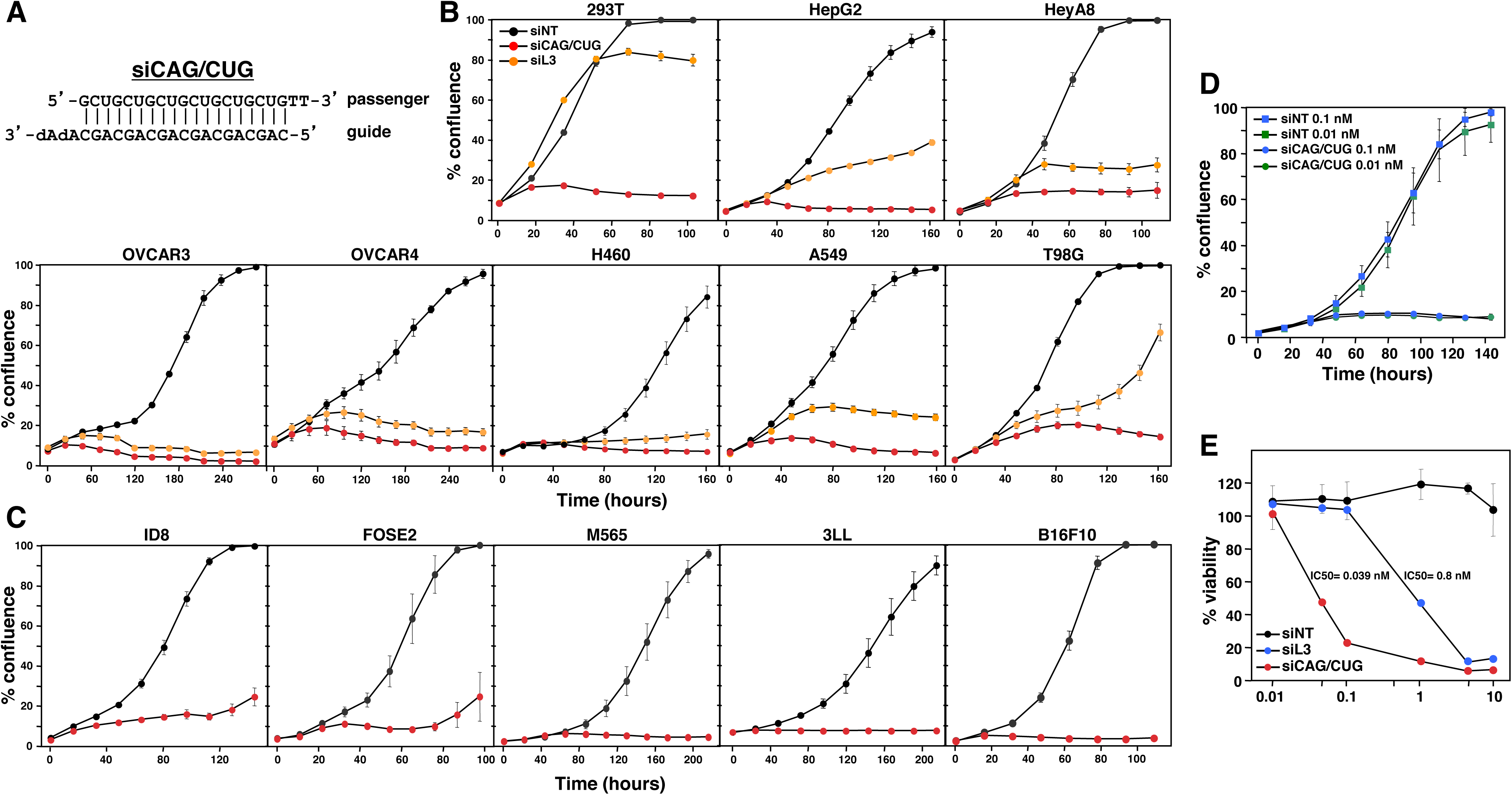
An siRNA duplex comprised of CAG and CUG repeats is super toxic to various cancer cell lines of human and mouse origin. **A** Sequence of the siCAG/CUG duplex. dA, deoxyadenosine. **B, C** Confluences over time of human (B) and mouse (C) cell lines transfected with 10 nM of either siNT, siL3, or siCAG/CUG. M565, 3LL and B16F10 cells were reverse transfected. Values are mean −/+SEM. The 13 cell lines were tested between 1 and 6 times in each case with between 3 and 6 technical replicates. **D** Confluence over time of HCT116 cells transfected with either siNT or siCAG/CUG at 0.1 or 0.01 nM. Values are mean −/+ SEM. n = 2 biological replicates, 6 technical replicates each. **E** Viability (ATP content) of HeyA8 cells transfected with different concentrations of siNT, siL3 or siCAG/CUG. Values are mean −/+ SD. n = 3 biological replicates, 3 technical replicates each.

### Identification of the most toxic TNR based siRNAs

The siCAG/CUG repeat 19mer in all three frames showed roughly the same level of toxicity when transfected into HeyA8 cells (Appendix Fig S2). To test whether other TNR disease-derived sequences were toxic to cancer cells when introduced as siRNAs, the repeats siGAA/UUC (GAA is amplified in Friedreich’s ataxia [22]), siCGG/CCG (CGG found in fragile X tremor ataxia syndrome [FXTAS] and CCG found in Fragile XE mental retardation [FRAXE] [1]) were transfected into HeyA8 (ovarian) and A549 (lung) cancer cells (Fig 2A). In addition, siCGA/UCG was tested because it has the same base composition as the super toxic siCAG/CUG TNR. Interestingly, among the four tested TNR siRNA duplexes two were super toxic to both cell lines, and two showed no toxicity. Most remarkable was the observation that siCGA/UCG was among the nontoxic repeats. This finding pointed at a sequence specific mechanism behind this phenomenon rather than a response of the cells to dsRNA of a specific base composition.

**Figure 2.**
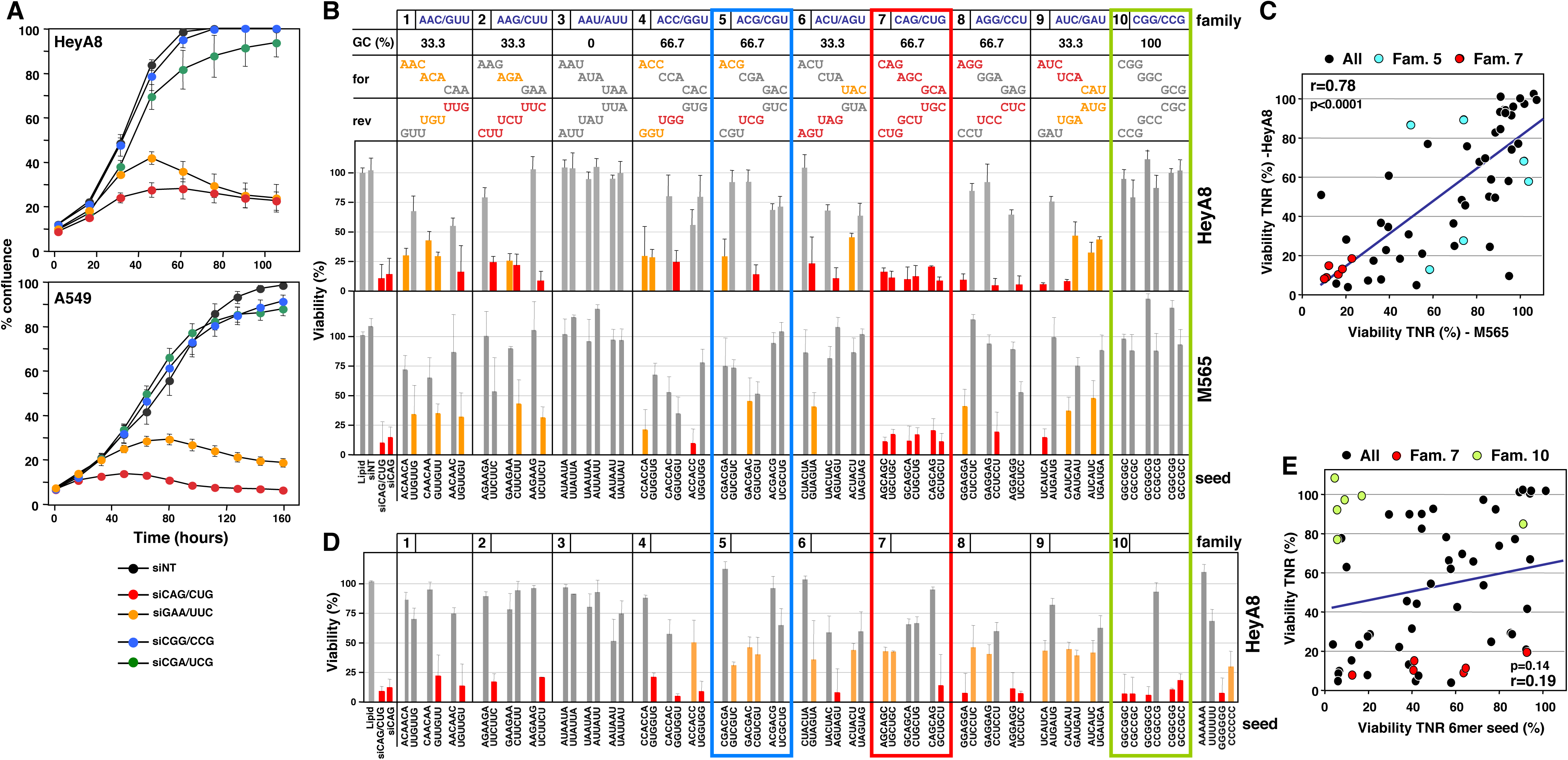
Identification of the most toxic TNR-based siRNAs. **A** Confluence over time of HeyA8 (top) or A549 (bottom) cells transfected with 1 nM of either siNT or the four TNR based siRNAs, siCAG/CUG, siCGG/CCG, siGAA/UUC and siCGA/UCG. Values are mean −/+SEM. n = 3/2 biological replicates (HeyA8/A549), 8 technical replicates. **B** Results of toxicity screens of 60 TNR-based siRNAs in human HeyA8 (top) and mouse M565 (bottom) cells. Cells were reverse transfected in triplicate in 384 well plates with 1 nM of each siRNA. In all cases, the complementary (sense) strand was inactivated by a 2’-OMe modification in positions 1 and 2. Grey: no or less than 50% loss in viability; Yellow: >50% loss in viability; red: >75% loss in viability. Family 7 is boxed in red as it is the only family in which all 6 duplexes were super toxic. Family 5 is boxed in blue. Its six members have the same GC content and the same nucleotide composition as the members of family 7. Family 10 is boxed in green. Each family contains three TNRs in the forward frame (for) and three TNRs that are the reverse complement (rev) of these TNRs. Values are mean −/+ SD. 3 technical replicates. **C** Correlation between the average of two viability screens performed in HeyA8 and two performed in M565 cells. The data points for the 6 members of TNR family 5 are labeled in blue those of family 7 are labeled in red. Pearson correlation and p-value is given. **D** Results of toxicity screen of 60 TNR-based 6mer seeds in a nontoxic backbone siRNA (see Appendix Fig S5B) in HeyA8 cells. Values are mean −/+ SD. n = 3 technical replicates. **E** Correlation between the average of two viability screens performed in HeyA8 with the 60 TNR based siRNAs and the screen performed with the 60 TNR based seeds. The data points for the 6 members of TNR family 7 are labeled in red those of family 10 are labeled in green. Pearson correlation and p-value is given.

To identify the most toxic TNR sequences in an unbiased screen, we designed a library of 19mer siRNAs based on the 60 possible TNRs (that contain more than one type of nucleotide). To reduce passenger strand loading and determine the toxicity of each repeat when loaded into the RISC as a guide strand, we replaced positions 1 and 2 of the passenger strand with 2’-O-methylated (OMe) nucleotides. To confirm the effect of the OMe modification, we modified the toxic CD95L-derived siRNA siL3 in this way. While the siL3 duplex modified on the intended passenger strand (S-OMe) was slightly more toxic to cells than unmodified siL3, likely reflecting a low level of passenger strand loading of siL3, neither siL3 modified on the antisense strand (AS-OMe) nor on both strands (S/AS-OMe) showed any toxicity (Appendix Fig S3).

All 60 TNRs were now synthesized with the sense strand carrying the OMe modification in positions 1 and 2, allowing us to determine the toxicity of each of the 60 antisense sequences. HeyA8 cells were transfected with 1 nM of each of the 60 TNRs and viability was quantified 96 hrs after transfection. The 60 TNRs can be grouped into 10 families [23]. Each family is comprised of 3 triplets shifted by one nucleotide plus its three complementary triplets. In total 30 (50%) of the TNRs were not toxic, 11 (18%) were moderately toxic (>50% loss of viability, shown in yellow), and 19 (31.7%) were super toxic (>75% loss of viability, shown in red) to HeyA8 cells (Fig 2B, top panels). Among the nontoxic TNRs were all 6 members of family 3 (0% GC content) and all 6 members of family 10 (100% GC content). All other TNR families contained nontoxic and toxic TNRs. Interestingly, in some cases just shifting the TNR sequence in the 19mer by one nucleotide resulted in opposite effects on viability (i.e. AGG and GGA in family 8). In other cases, members of a family showed toxicity of one strand but no toxicity of its complement (i.e. families 1, 2, 4, 5, 6, 8 and 9). This finding suggests a sequence-specific and in some cases frame specific activity of the TNRs consistent with RNAi being involved. Due to the different base composition of targeted RNAs, the comparison of TNR families with the same GC content and base composition is most meaningful. Two families contain a balanced GC content of 66.7% and identical base composition: family 5 and 7. Remarkably, while family 5 contained toxic and nontoxic members, all six TNRs in family 7 were super toxic (boxed in red in Fig 2B). Family 7 stands out as it contains all permutations of both the CAG and the CUG repeats we identified as killing all cancer cells.

To determine how much of these activities were conserved between human and mouse cancer cells, the screen was repeated with the mouse liver cancer cell line M565 (Fig 2B, bottom panels). The results for the siRNAs in TNR families 1, 2, 4, 5, 8, and 9 were somewhat similar to the ones obtained with the human cell line, but also showed clear differences. This could be due to differences in tissue origin, cell line, or species between the two cell lines. Three of the TNR families performed in an identical fashion between the two cell lines. Similar to HeyA8 cells, none of the 12 TNR-derived siRNAs in families 3 or 10 showed any toxicity in M565 cells. Most strikingly however, was the finding that again all 6 members of family 7, which contain both the CAG and the CUG repeat, were super toxic to the mouse cell line. Screens in both HeyA8 and M565 cells were repeated and results showed a high degree of congruence, especially in the results of family 7 (Appendix Fig S4). When the average of the screen in HeyA8 cells was plotted against the averages of the two screens in M565 cells, a significant correlation between the screens was found (Fig 2C) and again the six TNRs in family 7 were most consistently toxic. The data suggests the toxicity of this TNR family is conserved and it is independent of tissue, cell line and species.

We recently reported that the 6mer seed sequence of siL3 was the main determinant of its toxicity [15]. We therefore wondered how much of the toxicity of the super toxic TNRs was due to complete complementarity of the siRNA and how much was dependent on just the 6mer seed sequence. The data on siL3 were obtained by generating chimeric siRNA duplexes between a nontoxic control siRNA (siNT) and siL3 by replacing siL3 sequences from either end of the duplex with siNT sequences [15]. To generate an artificial nontoxic siRNA backbone in which to test all 60 TNR 6mer seed sequences, we first replaced 4 positions in the center of siNT still identical to the same position in the siL3 sequence with the complementary nucleotides, thereby removing any identity between siNT and siL3 outside the seed, while maintaining GC content (Appendix Fig S5A). This siL3 seed siRNA (siL3 seed) was almost as toxic to HeyA8 cells as siL3, confirming that the 6mer seed determined a substantial part of the toxicity of siL3. We therefore used the modified siNT backbone to test all possible TNR-derived 6mer seed sequences (Fig 2D, Appendix Fig S5B). While some TNR derived seeds were toxic to HeyA8 cells, there was only a moderate level of congruence between the screen with the entire TNR19mers and one just with the 6mers in the modified siNT backbone (Fig 2E). Of the 6 super toxic TNRs in family 7 only one was also toxic in the 6mer screen (Fig 2D and 2E). Interestingly, most of the 6mers in family 10 were toxic although no toxicity was observed in the TNR screen (Fig 2B). We interpret this as the inability of these 6 TNRs with their 100% GC content to properly enter the RISC. Together these data suggest 19mer TNR siRNAs are toxic to cancer cells by a mechanism distinct from the process of DISE which relies on just the seed sequences targeting the 3’UTRs of survival genes [15].

### Super toxic TNR-based siRNAs kill cancer cells through RNAi resulting in the loss of survival genes

To address the question whether the super toxic TNR-based siRNAs killed cancer cells through RNAi, we first compared the toxicity of siCAG/CUG in HCT116 wild-type and HCT116 Drosha^-/-^ cells. DISE inducing si- and shRNAs kill Drosha^-/-^ cells more efficiently than wild-type cells [15]. We had interpreted this as the RISC being more available in the absence of most cellular miRNAs, which rely on Drosha for processing. While siCAG/CUG was highly toxic to both cell lines at early time points, Drosha^-/-^ cells were more sensitive to growth reduction induced by siCAG than their wild-type counterparts (Appendix Fig S6, p=0.038, according to polynomial fitting model). To directly test the requirement of AGO2 in the siCAG/CUG induced toxicity we knocked down AGO2 in both HeyA8 and A549 cells (Fig 3A) and transfected the cells with either siNT or siCAG/CUG (Fig 3B). Removal of AGO2 from the cells almost completely prevented the toxicity of siCAG/CUG confirming dependence on the RISC. A dependence on Ago2 for siCAG/CUG toxicity was confirmed in *Ago1-4* knock-out mouse embryonic fibroblasts with re-expressed AGO2 (Appendix Fig. S7). These data indicated that siCAG/CUG was negatively affecting cells through canonical RNAi involving the RISC complex. To confirm this, we modified the siCAG siRNAs with the 2’-O-methylation to selectively block loading of either the siCAG or the siCUG based strand into the RISC (Fig 3C). When the CAG-based guide strand was modified (siCAG AS-OMe), the toxicity of the siCAG/CUG duplex was severely reduced. It was not affected when the CUG repeat containing strand was 2’-O-methylated (siCAG S-OMe), confirming that most of the toxicity of the siCAG/CUG repeat comes from the CAG repeat strand. siCAG/CUG did not have any toxicity when both strands were modified indicating most, if not all, of its toxicity requires RISC loading confirming that RNAi was responsible for cell death.

**Figure 3.**
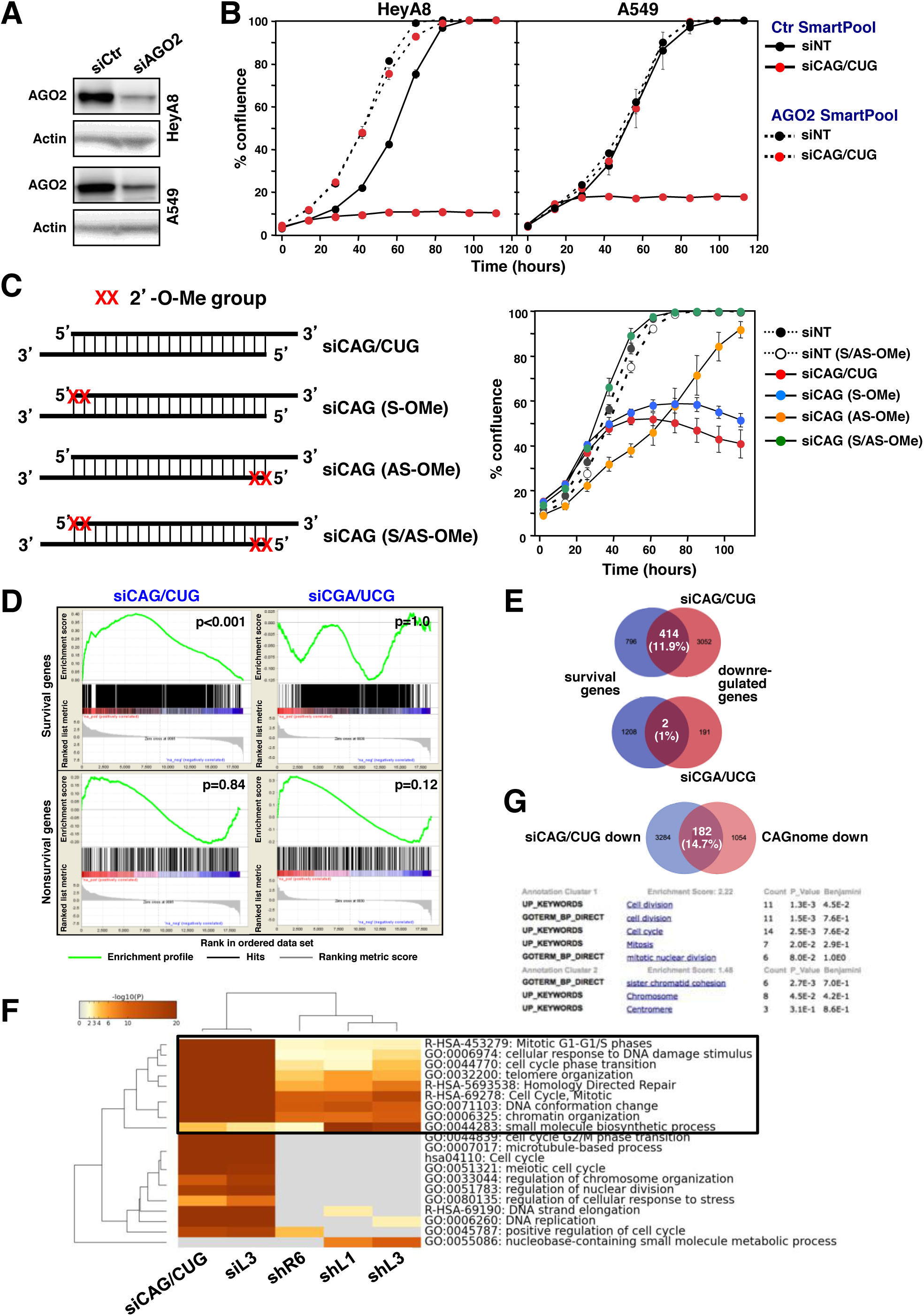
siCAG/CUG kills cancer cells through RNAi. **A** Western blot analysis of HeyA8 and A549 cells treated with a control SmartPool (siCtr) or an AGO2 siRNA SmartPool for 48 hrs. **B** Confluence over time of HeyA8 and A549 cells after transfection with 1 nM siNT or siCAG/CUG. Cells were first transfected with either 25 nM of a Ctr SmartPool or the AGO2 SmartPool and then after 24 hrs transfected with siNT or siCAG/CUG. Values are mean −/+SEM. n = 3 biological replicates, 3 technical replicates each. **C** *Left*: Sequences with positions of the 2’-O-methylation labeled in either the passenger/sense (S), the guide/antisense (AS) or both (A/AS) strands of siCAG/CUG. *Right*: Confluence over time of HeyA8 cells transfected with 10 nM of the four duplexes depicted on the left and two similarly modified duplexes derived from siNT. Values are mean −/+SEM. n = 2 biological replicates, 4-6 technical replicates each. **D** Gene set enrichment analysis for a group of 1846 survival genes (*top 2 panels*) and 416 nonsurvival genes (*bottom 2 panels*) identified in a genome-wide CRISPR lethality screen [45] after transfecting cells with either siCAG/CUG (left panels) or siCGA/UCG (right panels). Non targeting siNT served as control. p-values indicate the significance of enrichment. p values were calculated using the Kolmogorov-Smirnov test. **E** Venn diagrams showing the overlap between 1185 survival genes (genes identified as critical survival genes in two genome-wide lethality screens [45, 46], blue) and genes significantly downregulated (red) in cells transfected with either siCAG/CUG (top) or siCGA/UCG (bottom) when compared to siNT. **F** Metascape analysis of 4 RNA-Seq data sets of cells with introduced siL3, two CD95L derived shRNAs (shL1 and shL3), a CD95 derived shRNA (shR6) ORF, previously described [15], and the downregulated genes in cells transfected with siCAG/CUG. The boxed GO term clusters were highly enriched in all 5 data sets. **G** *Top,* Venn diagram comparing the 3466 genes downregulated in the siCAG/CUG treated HeyA8 cells(>1.5 fold adj. p <0.05) as determined by RNA-Seq (siCAG/CUG down) (see Fig 3D) and the 1236 genes expression of which inversely correlate with the length of CAG repeats recently reported in HD patients (CAGnome down) [24]. *Bottom,* DAVID gene ontology analysis of the overlapping 182 genes.

We recently reported that DISE-inducing CD95L derived sh- and siRNAs kill cancer cells by targeting the 3’UTR of critical survival genes through canonical RNAi [15]. To test whether the super toxic siCAG/CUG duplex also killed cancer cells through this mechanism, we transfected HeyA8 cells with siNT, siCAG/CUG or the nontoxic siCGA/UCG, and subjected the RNA 48 hours after transfection to a RNA-Seq analysis. Interestingly, in the cells transfected with siCAG/CUG, 3466 genes were down and 867 genes were upregulated (>1.5 fold, adjusted p value<0.05) (**Dataset EV1**). A DAVID gene ontology analysis of the upregulated genes did not reveal any evidence of an interferon response by the cells induced by the transfected siRNA (data not shown). In cells transfected with the nontoxic siCGA/UCG, only 194 genes were found to be downregulated and 420 genes upregulated.

Similar to cells undergoing DISE, when transfected with siCAG/CUG, ~1800 critical survival genes but not ~400 nonsurvival control genes [15] were significantly enriched in the downregulated genes in cells transfected with siCAG/CUG but not with siCGA/UCG in a gene set enrichment analysis (Fig 3D). In fact, we detected a ~12-fold increased percentage of survival genes compared to the non-survival genes among the downregulated RNAs in the siCAG/CUG treated cells (Fig 3E) - a higher difference than seen in cells treated with DISE-inducing sh- or siRNAs (data not shown). A Metascape gene ontology analysis comparing the downregulated genes in cells treated with either CD95 or CD95L derived si- or shRNAs with the data from the siCAG/CUG treated cells showed a strong overlap in the GO terms including cell cycle, response to DNA damage, mitosis, and chromatin organization suggesting that cells died through a mechanism similar to DISE (Fig 3F and Appendix Fig S8). When the RNA-Seq data of cells treated with siCGA/UCG was included in the analysis, not a single GO cluster overlapped (data not shown).

Interestingly, a large genome-wide comparison of lymphoblastoid cell lines from 107 HD patients reported an inverse correlation between CAG repeat length and downregulated genes. Biological pathways that were significantly affected were ribosomal process, energy metabolism and cell death pathways [24] all consistent with reduced cell viability. We compared the genes that were reported to be negatively and significantly correlated with the length of the CAG repeats in these patients (1236 genes, according to Pearson correlation) with the 3466 genes downregulated in the HeyA8 cells transfected with siCAG/CUG. Of the 1236 genes downregulated in the patients 182 (14.7%) were also downregulated in the siCAG/CUG treated HeyA8 cells (Fig 3G, top panel). In a DAVID gene ontology analysis with these 182 genes the two most significantly enriched clusters were consistent with genes playing a role in cell division and mitosis, consistent with a major effect of siCAG/CUG on mitosis (Fig 3G, bottom panel). In summary, these data suggest that the toxicity of the CAG repeat based siRNA may involve loss of survival genes and that this form of cell death could be related to the TNR activities seen in patients with extended CAG repeats.

### Super toxic TNR-derived siRNAs kill cells by targeting TNR sequences present in the ORF of genes complementary to the toxic siRNA guide strand

To determine which genes and what part of the mRNAs could be targeted by toxic TNR-derived siRNAs, we subjected ranked lists of downregulated genes of cells treated with either siCAG/CUG or siCGA/UCG to a Sylamer analysis [25]. This method detects enrichment of seed matches in mRNAs that are complementary to the seed of the introduced siRNAs. When the seed length was set to 6nts, we detected a minor enrichment of the 6mer TGCTGC in the 3’UTRs of the downregulated genes in the cells treated with siCAG/CUG (Appendix Fig S9A, left panel). TGCTGC is the expected seed match (position 2-7) of the siCAG 19mer guide strand. No significant seed match enrichment was found in cells treated with siCGA/UCG or when the ORFs of these genes were analyzed (Appendix Fig S9A, right panel and S9B). However, when we analyzed the ORFs of cells treated with siCAG/CUG and even when setting the seed length to the maximum of 10 nts, we found a very profound enrichment of two 10 nt sequences (p-value ~10^-50^) that corresponded to positions 1-10 and 2-11, respectively of the targeting siCAG 19mer (Fig 4A). No such enrichment was found when the 3’UTRs of the genes were used for the analysis (Appendix Fig S9C). These data suggest that in contrast to DISE-inducing si/shRNAs, siCAG/CUG killed cancer cells by targeting long repeat sequences located mainly in ORFs. Consistent with this conclusion, genes containing either of the two targeted 10mers in their ORFs were very strongly enriched among the downregulated genes in siCAG/CUG treated cells, while only a weak enrichment was found when the 3’UTRs were analyzed (Fig 4B).

**Figure 4.**
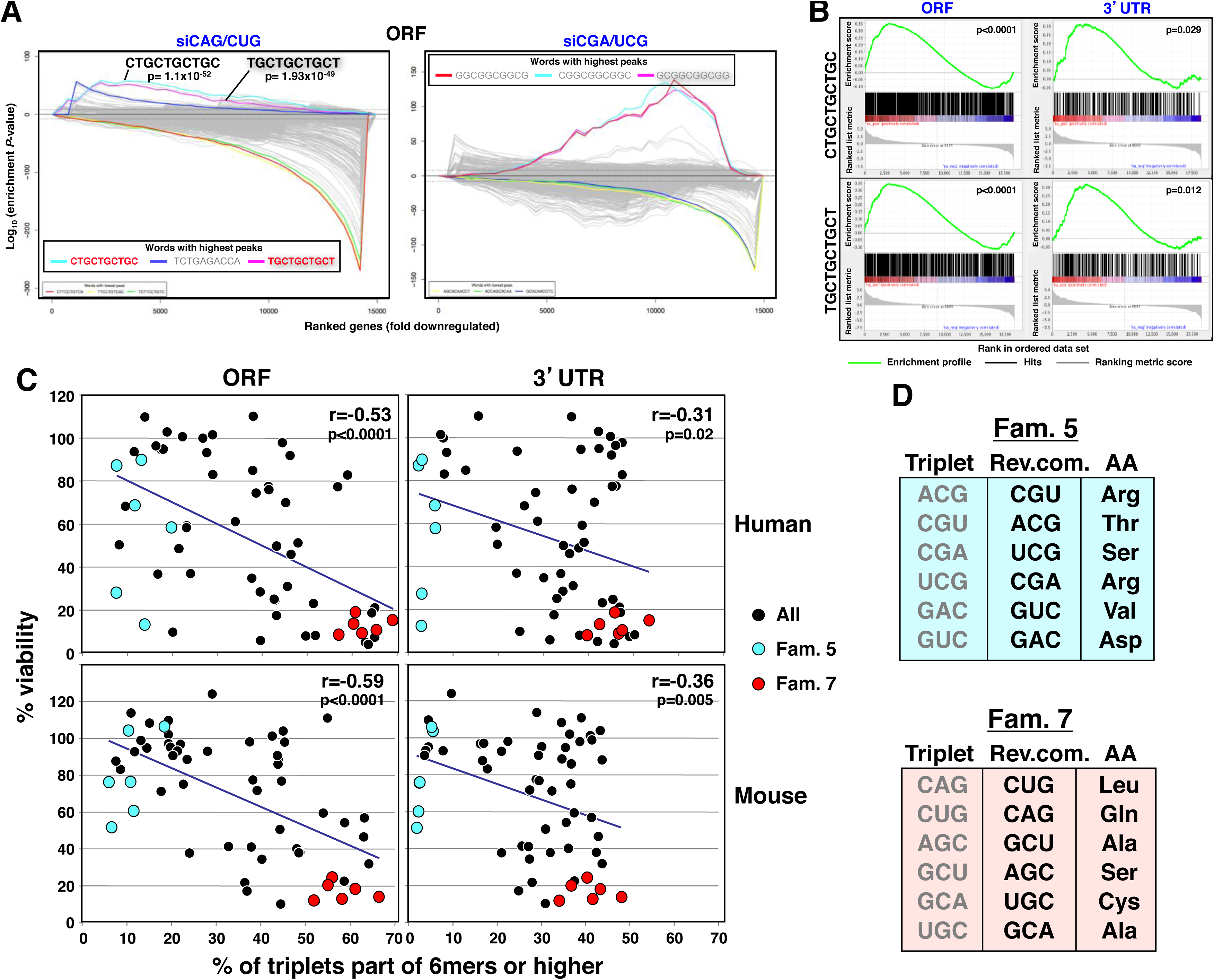
The toxicity of the TNR-based siRNAs correlates with the presence of long complementary repeats in the ORFs of genes. **A** 10 nucleotide Sylamer analysis of the ORFs of genes deregulated in HeyA8 cells transfected with either siCAG/CUG (left) or siCGA/UCG (right) ordered from most down-regulated to most up-regulated. The three most highly enriched sequences are shown. The 10 nucleotide motifs in bold in the siCAG/CUG analysis correspond to the reverse complement of position 1-10 and 2-11 in the 19mer siCAG strand. Bonferroni-adjusted p-values are shown. Sequences without a p-value were either not significantly enriched or not at the top of the ranked list. **B** Gene set enrichment analysis for genes containing the CTGCTGCTGC (*top panels*) or TGCTGCTGCT (*bottom panels*) sequences identified as enriched in the downregulated genes (see A) in cells transfected with siCAG/CUG when compared to cells transfected with siNT. The analyses were performed with either the ORF (*left panels*) or the 3’UTR (*right panels*) of the downregulated genes. p-values indicate the significance of enrichment. p values were calculated using the Kolmogorov-Smirnov test. **C** Correlation between the average of the two viability screens performed in HeyA8 (*top panels*) or M565 (*bottom panels*) cells with the percentage of TNRs that are part of 6mer or higher longer repeat sequences in either the ORF (*left panels*) or 3’UTR (*right panel*) of the genes. The data points for the 6 members of TNR family 5 are labeled in blue and those of family 7 are labeled in red. Pearson correlations and p-values are given. **D** The amino acids that are coded by the targeted triplets in families 5 and 7.

Now knowing that the toxicity of the siCAG/CUG correlated with the presence of targeted trinucleotide repeats found in the ORFs of genes, we wondered whether the toxicity across all 60 TNR derived siRNAs correlated with the presence of higher ordered reverse complementary TNRs in ORFs that could be targeted by the TNR siRNAs and whether this was conserved between human and mouse cells. When counting all 60 single triplets in both the ORFs and 3’UTRs of all human and mouse genes they were found to be slightly more abundant in the ORF of genes (Appendix Fig S10A). When separating into individual triplets of the 10 families most triplets are found at similar frequencies in both the ORFs and 3’UTRs in humans and in mice (Appendix Fig S10B, top panels in each set). This situation changed when we focused on targeted (reverse complements of the targeting TNRs) repeats. We plotted the results for 6mers (as this is the minimum sequence required for RNAi-based targeting), 10mers (the maximum seed length allowed by Sylamer), and 19mers (the length of the siRNAs used). In all cases, with longer sequences, the preference for certain triplets became clearer (Appendix Fig S10B, bottom three panels in each set). When analyzing the 19mers, in both human and mouse, the most abundant TNRs are members of the super toxic family 7 and barely any 19mers were found in the related family 5. Genes containing the 19mer TNR (or longer) in their ORF targeted by siCAG were enriched in the most downregulated genes in cells transfected with siCAG/CUG (Appendix Fig S10C). We selected 5 of the 6 most highly expressed and downregulated genes from the RNA-Seq analysis for validation (**Dataset EV2 and** Fig S11). HeyA8 cells were transfected with siRNAs at 1 nM, and the mRNAs levels were quantified by real time PCR 10, 20 or 40 hrs after transfection. When transfecting siCAG/CUG all 5 CUG repeat-containing mRNAs were downregulated as early as 10 hrs with maximal downregulation at 40 hrs. Specificity of the targeting was established by transfecting the cells with either siCAG or siCUG (in which the passenger strand was disabled by adding the 2’-O-methylation). Only the siCAG-based siRNAs were active in silencing the CUG TNR containing genes. These data strongly suggested that the toxicity of TNR-based siRNAs in general might be explained by the presence of extended reverse complementary repeat sequences present preferentially in the ORF of targeted genes. This was confirmed by plotting the viability of cells treated with any of the 60 TNR siRNAs and the number of targeted TNR sequences of 6nts or longer (Fig 4C). The highest and most significant correlation was found for both human and mouse ORFs. Remarkably, while 80-90% of triplets targeted by the 6 members of the CGA containing family 5 are present as singular events (blue dots), between 50 and 70 % of the triplets targeted by members of the CAG containing family 7 are found to be part of 6mers or higher ordered TNRs (red dots). Interestingly, the TNRs targeted by the 6 toxic members of family 7 code for 5 different amino acids (Fig 4D). As all the 6 members of family 7 are equally toxic to cancer cells, this suggests that this involves targeting long repeat elements (i.e. TNRs), rather than a requirement for poly-homo amino acid coding stretches. This view is supported by an analysis of species conservation (Appendix Fig S12). Of the genes that contain targeted 19mers, only 7 of the 99 genes found in either the mouse and the human genome overlapped and the genes that are targeted do not have shared functions (Appendix Fig S12B). In summary, our data provide evidence that TNR-based siRNAs are toxic to cells by targeting a number of genes that contain high order trinucleotide repeats that are reverse complementary to the targeting TNR. The resulting cell death has features of what we recently described as DISE, with the main difference that DISE is the result of a miRNA-like targeting of short seed matches in the 3’UTRs of survival genes, whereas the TNR-induced cell death is an on-target effect affecting a larger number of genes that contain targeted sequences in their ORF. This now provides an explanation for why the most toxic TNR-based siRNAs are much more toxic than DISE-inducing siRNAs. Intriguingly, the CAG repeats found in HD are part of the most toxic family of TNRs and their reverse complementary 19mers that can serve as targets are the most abundant TNR sequences in the ORFs of both human and mouse genomes.

### Super toxic CAG/CUG TNR based siRNAs slow down tumor growth *in vivo* with no toxicity to normal tissues

We were wondering whether the super toxic TNR-based siRNAs could be used for cancer therapy. We decided to deliver the siRNAs to cancer cells *in vivo* using templated lipoprotein (TLP) nanoparticles [26]. Before using the TLP particles loaded with the siCAG/CUG duplex (siCAG/CUG-TLP) *in vivo,* we tested their effects on tumor cells *in vitro.* They killed HeyA8 cells more efficiently than siL3-TLPs (Fig 5A) and also slowed down growth of the tested human or mouse cancer cell lines (Fig 5B and data not shown). They also killed neurospheres derived from patients with glioblastoma (Fig 5C). To test the activity of the siCAG/CUG-TLPs *in vivo* we treated orthotopically xenografted HeyA8 ovarian cancer cells in mice. Mice injected with 100,000 tumor cells were i.p. injected with nano particles 5 times a week for two weeks (Fig 5D). After the tenth treatment mice were split into two groups, one group continued to receive treatment in the third week and the second group did not. This was done to determine whether large established tumors would still respond to the treatment. The large tumors in treatment group 1 still benefited from the effect of siCAG/CUG in the third week of treatment (Fig. 5E, left panel). In contrast, some tumors in the mice in treatment group 2 grew out rapidly, while others showed persisting growth reduction (Fig. 5E, right panel). These results suggest that established tumors respond to the siCAG/CUG treatment. This was confirmed in another experiment in which 10^6^ HeyA8 cells were injected and mice were first treated 3 times a week and then switched to daily treatment 19 days after tumor cell injection (data not shown).

**Figure 5.**
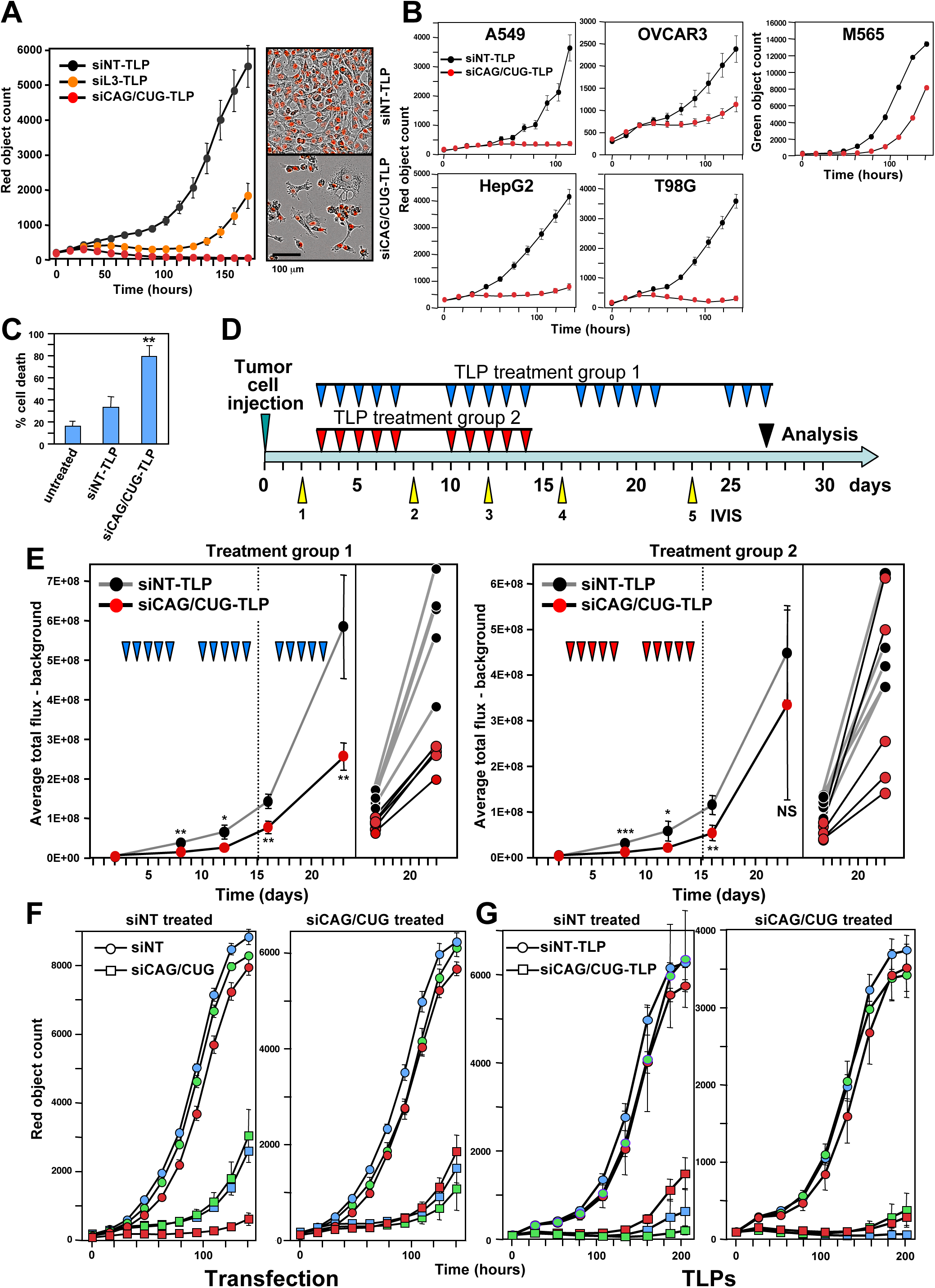
Killing cancer cells using siCAG/CUG coupled to TLP nanoparticles both *in vitro* and *in vivo*. **A** *Left:* Confluency over time of HeyA8 (Nuc-Red) cells treated with 10 nM of either siNT-TLP siL3-TLP, or siCAG/CUG-TLP. *Right*: Phase and red fluorescence image of HeyA8 (Nuc-Red) cells 90 hrs after transfection with 15 nM of TLPs. Values are mean −/+SEM. n = 3 technical replicates. **B** Confluency over time of different human and mouse cancer cell lines treated with either 10 nM (OVCAR3, HepG2, M565) or 20 nM (A549, T98G) of TLPs. Values are mean −/+SEM. n = 2 biological replicates, 6 technical replicates each. **C** Percent cell death (Trypan blue counting) of GIC-20 neurospheres derived from a patient with glioblastoma six days after adding the TLPs (30 nM). Values are mean −/+SD. n = 3 technical replicates. **p<0.01. **D** Treatment scheme. **E** Tumor growth over time based on small animal imaging of 10^5^ HeyA8-Nuc-red-Luc-neo cells injected i.p. into NSG mice treated with either siNT-TLPs or siCAG/CUG-TLPs. Treatment group 1 received 18 injections over four weeks and treatment group 2 10 injections over three weeks. The bioluminescence signal of IVIS #4 and #5 for individual mice is shown (right panel). The experiment represents one of two similar experiments. Values are mean −/+SD. * p<0.05; **p<0.01; ***p<0.0001, NS, not significant. **F, G** Change in red object count (growth) of tumor cells from 3 mice of the siNT-TLP and the siCAG/CUG-TLP treatment group 1 either after transfection with 1 nM siNT or siCAG/CUG (F) or after incubation with 7.5 nM siNT-TLP or siCAG/CUG-TLP (G). 1000 cells per well were plated. Values are mean −/+SEM. n = 3 - 8 technical replicates.

To determine whether siCAG/CUG was detrimental to mice, mice in treatment group 1 were treated a few more times with the siRNA and were analyzed just when the control treated mice were moribund at around day 27. We did not see any signs of toxicity in any of the mice. They were feeding well (not shown), did not lose weight (Appendix Fig S13A), had normal liver histology (Appendix Fig S13B), and showed no increase in liver enzymes in the serum (Appendix Fig S13C). These data demonstrated that super toxic CAG/CUG TNR-based siRNAs delayed tumor growth *in vivo* 5-6 days with no gross toxicity to normal cells and that they could be safely administered using TLP nanoparticles.

To determine whether tumor cells acquired resistance to the treatment, we tested tumors from three mice in treatment group 1 *ex vivo*. Three tumors of mice treated with siNT-TLP and three tumors of mice treated with siCAG/CUG-TLP were transfected with the same siRNAs *in vitro* a day after tumor isolation (Fig 5F). In parallel, these tumor cells were also treated with the nanoparticles again (Fig 5G). In all cases, the tumors from the mice that had received the toxic siRNA were as sensitive to the toxic effects of siCAG/CUG *in vitro* as the tumors from mice treated with siNT suggesting that cancer cells cannot become resistant to cell death induced by the toxic TNR-based siRNAs, at least not in the classical sense observed after targeted therapy, and that preferentially tumor cells were responding as there were no signs of toxicity in the mice.

## Discussion

The siRNA we used for cancer treatment was a duplex between the basic TNR module found in HD (CAG) and the fully complementary strand CUG found in Myotonic Dystrophy (DM1). In a screen of all TNR-derived siRNAs both CAG and CUG were part of a family that contains 6 members all of which were highly toxic to cancer cells of both human and mouse origin. This hybrid duplex between the two disease molecules was also recently tested in a well-established *Drosophila* model of DM1 [27]. Expression of the two transcripts led to the generation of Dicer-2 (dcr-2) and ago2-dependent 21-nt TNR-derived siRNAs resulting in high toxicity to the cells. In a separate study, it was shown that expression of these complementary repeat RNAs leads to dcr-2-dependent neurodegeneration [28]. These results suggest that co-expression of CAG and CUG repeat-derived sequences may dramatically enhance toxicity in human repeat expansion diseases in which anti-sense transcription occurs. Antisense transcription was reported to occur in SCA8 in two genes encompassing the repeats are expressed: ATXN8 (CAG repeat), on the sense strand and ATXN8OS (CTG repeat) on the antisense strand [29].

One could argue that a cancer therapy based on delivering siCAG/CUG could be detrimental to patients as TNR expansion patients suffer from various pathologies. However, similar to many other genes with amplified CAG repeats, HTT is ubiquitously expressed throughout the body with somewhat higher expression in the brain and in testis [30]. The disease is characterized by neurodegeneration affecting the cerebral cortex and neuropathology in the striatum, but it also affects other tissues [31]. So, if sCAGs are produced in multiple tissues the effects on most normal tissues seems to be moderate. Even in the brain, while detrimental to HD patients long term, most patients do not have major symptoms before the age of 40 [4, 31]. Short-term exposure to toxic sCAGs for cancer therapy, as suggested by our mouse experiments, may not have a dramatic effect on normal tissues but may be enough to kill cancer cells. If a CAG based siRNA were to produce side effects particularly in the brain it may be possible to protect the brain through local administration of neuroprotecting LNA-CTGs as described [14].

What could be the mechanism of the relative resistance of normal versus tumor cells to the toxic siRNAs? Our recent data suggested that miRNAs inhibit DISE and in fact may protect normal cells from it [15]. Both Drosha and Dicer k.o. HCT116 cells were found to be hypersensitive to a DISE inducing siRNA. This is entirely consistent with reported activities of CAG repeats which may also act though RNAi. Pathogenic Ataxin-3 with amplified CAG repeats showed strongly enhanced toxicity in HeLa cells after knockdown of Dicer [32]. In addition, it was previously shown in *Drosophila* that impairing miRNA processing dramatically enhanced neurodegeneration caused by the CAG repeat gene Ataxin-3. Two fly mutants were tested, one with a deficiency in dcr-1, the Drosophila Dicer ortholog that is required for miRNA biogenesis, and another with a deficiency in R3D1, a gene required for dcr-1 to function [33]. The authors concluded that "miRNA pathways normally play a protective role in polyQ-induced neurodegeneration". In the light of our data it is possible that miRNAs might actually protect cells from the toxic effects of TNR based siRNAs.

While HD is the best known disease caused by CAG repeats, one of the two diseases first discovered to be caused by TNR repeat expansions is the neurodegenerative disorder SBMA/Kennedy disease [34] wherein the pathogenic CAG repeat is found in exon 1b of the androgen receptor (AR). If patients with amplified CAG repeats produce sCAGs, which according to our work may be detrimental to cancer cells, one would have to expect that such patients have a reduced cancer incidence due to toxic siRNA expression in many tissues. Indeed reduced cancer incidence was reported for HD and SBMA patient populations in Sweden [18], and HD patients in France [19], Denmark [20], and England [21].

Possibly the clearest connection to cancer has been reported for the CAG repeats in the AR gene and prostate cancer (PCa). The CAG repeat length in the AR has been inversely linked to PCa. While longer repeats (>20 CAGs) confer a protective effect among the PCa patients 45 years or older [35], shorter CAG repeats have been shown to result in a two-fold increased cancer risk [36], a more aggressive disease, and a high risk of distant metastases [37–39]. Shortening of CAG repeat length was found in *in situ* lesions of PCa and its possible precursors [40], suggesting that PCa avoids longer CAG repeats. This is consistent with our finding of super toxicity of CAG based siRNAs.

There are two observations that suggest the targeted TNRs present in the ORFs of certain genes are not there because these proteins require stretches of the same amino acid for their function which would presumably be conserved between human and mouse: first, all six members of the TNR family 7 were super toxic targeting 6 different reverse complementary TNRs that code for 5 different amino acids (Fig 4D), and second, the genes with the longest repeats with complete complementarity to the most toxic siCAG/CUG of 19 nts showed little overlap between human and mouse (Appendix Fig S12B) and the different targeted genes do not share similar functions (Appendix Fig S12C). Interestingly, both the AR and the HTT genes contain some of the longest CAG repeats in the human genome, however, those are not found in the mouse orthologs at the same positions in the ORF.

So, if there was no pressure to maintain these TNRs in specific genes but rather anywhere in the genome, could there be an evolutionary link between TNRs and cancer? A hint may come from the way the repeat expansion are generated. It is believed that among other mechanisms DNA replication slippage and/or defective base excision repair causes expansion of TNRs [41]. Therefore CAG repeats could be part of a mechanism used during evolution to maintain genome integrity and, in the context of multicellular organisms, to prevent cancer formation by producing toxic siRNAs. This would occur whenever too many mutations start accumulating in cells, one property all cancers have in common.

While the treatment with siCAG/CUG requires optimization, our data on the toxicity of CAG TNR based siRNAs for cancer cells but not normal cells when administered *in vivo* and the reported decreased incidence rate for different types of cancer in patients with CAG expansions suggest that TNR-based siRNAs may be useful for cancer therapy.

## Materials and Methods

### Cell lines and tissue culture

All cells were grown in an atmosphere of 5% carbon dioxide (CO_2_) at 37°C. Unless indicated otherwise base media were supplemented with 10% heat-inactivated fetal bovine serum (FBS; Sigma-Aldrich) and 1% penicillin/streptomycin and L-Glutamine (Mediatech Inc.). Cells were dissociated with 0.25% (w/v) Trypsin-0.53 mM EDTA solution (Mediatech Inc.). The following cell lines were cultured in supplemented RPMI1640 Medium (Mediatech Inc.): Ovarian cancer cell lines HeyA8 (RRID:CVCL_8878), OVCAR3 and OVCAR4 (both from Tumor Biology Core, Northwestern University), and lung cancer cells A549 (ATCC CRM-CCL- 185 and H460 (ATCC HTB-177). The GBM cell line T98G (ATCC CRL1690) was cultured in Eagle’s Minimum Essential Medium (EMEM) (ATCC). Melanoma B16F10 cells (ATCC CRL-6475) and 293T cells (RRID:CVCL_0063) were cultured in DMEM (Cellgro). HepG2 (ATCC HB-80645) was cultured in EMEM (ATCC). ID8, a mouse ovarian cancer cell line, was cultured in DMEM supplemented with 4% FBS, and 10 mg/l Insulin, 5.5 mg/l Transferrin, 6.7 µg/ml Selenium (ITS, Mediatech, Inc., 1:10 diluted). and 3LL Lewis lung cancer cells (ATCC CRL-1642) were cultured in DMEM. FOSE2 cells are spontaneously immortalized ovarian surface epithelial cells, and M565 cells are from a spontaneously formed liver cancer in a female and male mouse, respectively, both isolated from mice carrying a floxed Fas allele [17]. Both were cultured in DMEM/F12 (Gibco #11330), 1% ITS (Insulin-Transferrin-Selenium Gibco 51300-044). M565 cells were dissociated with Accutase detachment reagent (Fisher Sci.). HCT116 Drosha^**−/−**^ were generated by Narry Kim [42]. HCT116 parental (cat#HC19023, RRID:CVCL_0291) and the Drosha^-/-^ clone (clone #40, cat#HC19020) were purchased from Korean Collection for Type Cultures (KCTC). All HCT116 cells were cultured in McCoy’s medium (ATCC, cat#30-2007). Mouse *Ago1*-4 k.o. embryonic stem cells inducibly expressing human FLAG-HA-AGO2 were described in [43]. CELLSTAR tissue culture dishes (Greiner Bio-One, cat#664160, cat#639160) were coated with 0.1% Gelatin solution (Sigma, cat#ES-006-B) for 10 to 30 minutes before use. Cells were cultured in DMEM media (Gibco, cat#12430054) supplemented with 15% Fetal Bovine Serum (Sigma, cat#F2442), 1% NEAA solution (HyClone, cat#SH3023801), 1% GlutaMAX 100x (Gibco, cat#35050061), 0.0007% 2-Mercaptoethanol (Fisher, cat#BP176100), and 10^6^ units/L LIF (Sigma, #ESG1107). The cell culture media was refreshed daily. FLAG-HA-AGO2 is under the control of a TRE-Tight (TT) doxycycline (Dox)-inducible promoter. 100 ng/ml doxycycline (Sigma, cat#D9891) was added to the media in order to induce moderate level of Ago2 expression to maintain normal cell growth. To deplete Ago2 expression in cells, doxycycline was withdrawn from media for 4 days. To induce wild type level of hAgo2 expression, 2.5μg/ml doxycycline was added to the media. The human GBM derived neurosphere cell line GIC-20 (infected with pLV-Tomato-IRES-Luciferase) was obtained from Dr. Alexander Stegh. Cells were grown as neurospheres in DMEM/F12 50:50 with L-glutamine (Corning), supplemented with 1% PenStrep, B27 (Invitrogen), N2 (Invitrogen), human-Epidermal Growth Factor (hEGF; Shenandoah Biotech), Fibroblast Growth Factor (FGF; Shenandoah Biotech), Leukemia Inhibitory Factor (LIF; Shenandoah Biotech), and GlutaMAX (Life Technologies). HeyA8 xenografted tumors nodules were dissected from mice, cut, washed in sterile PBS, and dissociated with dissociated with 0.25% (w/v) Trypsin-0.53 mM EDTA solution for 20 minutes at 37°C. The digestions was stopped by adding full RPMI-1640 medium. After centrifugation, the trypsin solution mix was removed, and the tumor cells were resuspended in fresh full medium, and strained through 70 micron cell strainer. The tumor cell suspension was plated over night on 10 cm tissue culture dishes. The following day, cells were harvested, counted, and plated on 96-well plates for further experiments. Cells were transfected with siRNAs after cells had adhered or incubated with siRNA TLPs and then plated.

### Western blot analysis

Primary antibodies for Western blot: anti-β-actin antibody (Santa Cruz #sc-47778, RRID:AB_626632), anti-human AGO2 (Abcam #AB186733, RRID:AB_2713978). Secondary antibodies for Western blot: Goat anti-rabbit; IgG-HRP (Southern Biotech #SB-4030-05, RRID:AB_2687483). Reagents used: propidium iodide (Sigma-Aldrich #P4864), puromycin (Sigma-Aldrich #P9620) and Lipofectamine RNAiMAX (ThermoFisher Scientific #13778150). Western blot analysis was performed as recently described [15]

### Transfection with short oligonucleotides

For transfection of cancer cells with siRNAs, RNAiMAX was used at a concentration optimized for each cell line, following the instructions of the vendor. Cell lines were either transfected after cells had adhered (forward transfection), or during plating (reverse transfection). For an IncuCyte experiment cells were typically plated in 200 μl antibiotic free medium, and 50 μl transfection mix with RNAiMAX and siRNAs were added. During growth curve acquisitions the medium was not exchanged to avoid perturbations. All individual siRNA oligonucleotides were ordered from Integrated DNA Technologies (IDT). Individual RNA oligos were ordered for the sense and antisense oligo; the sense strand had 2 Ts added to the 3’ end; antisense strand had 2 deoxy As at the 3’ end. When indicated the first two positions at the 5’-end were 2’-O-methylated. Sense and antisense oligos were mixed with nuclease free Duplex buffer (IDT, Cat.No# 11-01-03-01; 100 mM Potassium Acetate, 30 mM HEPES, pH 7.5) to 20 μM (working solution), heated up for 2 minutes at 94°C, and then the oligos were allowed to cool down to room temperature for 30 minutes. siRNA solutions were aliquoted and stored at -80°C. The cells were transfected with siRNAs at a final concentration of 0.01 nM - 10 nM. The following siRNA sequences were used: siNT (siNT#2): UGGUUUACAUGUUGUGUGA (non targeting in mammalian cells), siNT1: UGGUUUACAUGUCGACUAA (non targeting in mammalian cells), siL3: GCCCUUCAAUUACCCAUAU (human CD95L exon 1), siNT/siL3: UGGUUUACAUGUCCCAUAA. siNT seed: UGGUAAACUAGUUGUCUGA, siL3 seed: UGGUAAACUAGUCCCAUAA. All TNR based 19mer siRNAs were designed as follows: The TNR based siRNA was named according to its antisense/guide strand: 2nt 3’ overhangs were added as described above. All TNR based siRNAs were fully complementary 19mers. For instance, the siCAG/CUG sequences are: S: CAGCAGCAGCAGCAGCAGCTT, AS: GCUGCUGCUGCUGCUGCUGTT. siRNA duplexes used in the screens are shown in **Dataset EV3**. In all siRNAs used in screens the sense/passenger strand was disabled by 2’-O-methylation in positions 1 and 2 of the sense strand.

For transfecting *Ago1-4* k.o. mouse ESC, cells were cultured without doxycycline for three days. Half of the cells were then cultured for one more day without doxycycline before transfection. The other half was cultured in media containing 2.5 µg /ml of doxycycline for one day to induce WT level AGO2 expression before transfection. For IncuCyte experiments, The ESCs were transfected with either 5 nM siNT or siCAG/CUG using reverse transfection method in a 96 well plate coated with 0.1% gelatin. 5000 cells/well and 0.2 µl RNAiMAX/well were used. One day after transfection, 100 µl of media (with or without 2.5 µg/ml doxycycline) was added to each well. After that, media was refreshed every 2~3 days until the cells grew confluent. For flow cytometry experiments, cells were transfected with either 5 nM unlabeled siNT or siNT labeled with Cy5 on the 5’ end of the antisense strand using reverse transfection method in a 12 well plate coated with gelatin in triplicates. 300,000 cells/well and 1 µl RNAiMAX/well were used. Flow cytometry measurements were conducted 24 hours after transfection. For AGO2 knockdown experiment, 100,000 cells/well HeyA8 or 200,000 cells/well A549 cells were reverse transfected in 6-well plate with either non-targeting (Dharmacon, cat#D-001810-10-05) or an AGO2 targeting siRNA SMARTpool (Dharmacon, cat#L004639-00-005) at 25 nM. 1 µl RNAiMAX per well was used for HeyA8 cells and 6 µl RNAiMAX per well was used for A549 cells. 24 hours after transfection with the SMARTpools, cells were reversed transfected in 96-well plate with either siNT or siCAG/CUG at 1 nM and monitored in the IncuCyte. 0.1 µl/well RNAiMAX was used for HeyA8 cells and 0.6 µl/well RNAiMAX was used for A549 cells.

### Total RNA isolation and RNA-seq analysis

HeyA8 cells were transfected in 6-wells with siNT or either siCAG/CUG or siCGA/UCG oligonucleotides at 1 nM. The transfection mix was removed after 9 hours. Total RNA was isolated 48 hours after transfection using the miRNeasy Mini Kit (Qiagen, Cat.No. 74004) following the manufacturers instructions. An on-column digestion step using the RNAse-free DNAse Set (Qiagen, Cat.No.: 79254) was included. NGS RNA-SEQ library construction and sequencing was performed by the University of Chicago Genomics Facility. The quality and quantity of RNA samples was assessed using an Agilent bio-analyzer. RNA-SEQ libraries were generated using Illumina Stranded TotalRNA TruSeq kits according to the Illumina provided protocol and sequencing was performed using the Illumina HiSEQ4000 according to Illumina provided protocols and reagents. The resulting paired end reads were aligned to the hg38 assembly of the human genome with Tophat2. HTseq was used to associate the aligned reads with genes, and EdgeR was used to identify genes significantly differentially expressed between treatments, all as recently described [15]. The accession number for the RNA-Seq and expression data reported in this work are GSE104552.

### Real-time PCR

Real-time PCR was performed a described recently [15] using the following primers: GAPDH (Hs00266705_g1), RPL14 (Hs03004339_g1), LRRC59 (Hs00372611_m1), CNPY3 (Hs01047697_m1), CTSA (Hs00264902_m1), and LRP8 (Hs00182998_m1).

### Monitoring growth over time and quantification of cell death

To monitor cell growth over time, cells were seeded between 125 and 4000 per well in a 96-well plate in triplicates. The plate was then scanned using the IncuCyte ZOOM live cell imaging system (Essen BioScience). Images were captured at regular intervals, at the indicated time points, using a 10x objective. Cell confluence was calculated using the IncuCyte ZOOM software (version 2015A). IC50 values for siL3 and siCAG/CUG were determined using GraphPad Prism 6 software (by logarithm normalized sigmoidal dose curve fitting). Quantification of DNA fragmentation (subG1 DNA) was done as previously described [15].

### siRNA screens and cell viability assay

HeyA8 or M565 cells were expanded and frozen down at the same passage. One week before transfection, cells were thawed and cultured in RPMI1640 medium, 10% FBS and 1% pen/strep. Cells were split three times during the week and each time seeded at 4x10^6^ cells total in one T75 flask. On the day of the transfection, RNA duplexes were first diluted with Opti-MEM to make 30 μl solution of 10 nM (for the duplexes with the 6mer seeds) or 1 nM (for the TNR-based duplexes) as final concentration in a 384-well plate by Multidrop Combi. Lipofectamine RNAiMAX (Invitrogen) was diluted in Opti-MEM (6 μl lipid + 994 μl of Opti-MEM for HeyA8 and 15.2 μl lipid + 984.8 μl of Opti-MEM for M565 cells). After incubating at room temperature for 5 to 10 minutes, 30 μl of the diluted lipid was dispensed into each well of the plate that contains RNA duplexes. The mixture was pipetted up and down three times by PerkinElmer EP3, incubated at room temperature for at least 20 minutes, and then the mixture was mixed again by PerkinElmer EP3. 15 μl of the mixture was then transferred into wells of three new plates (triplicates) using the PerkinElmer EP3. 50 μl with 320 HeyA8 or 820 M565 cells was then added to each well containing the duplex and lipid mix, which results in a final volume of 65 μl. Plates were left at room temperature for 30 minutes then moved to a 37^o^C incubator. 96 hours post transfection, cell viability was assayed using CellTiter-Glo (Promega) quantifying cellular ATP content. 35 μl medium was removed from each well and 30 μl CellTiter-Glo cell viability reagent was added. The plates were shaken for 5 minutes and incubated at room temperature for 15 minutes. Luminescence was then read on the BioTek Synergy NEO2.

### Treatment of xenografted ovarian cancer cells *in vivo* with templated lipoprotein particles (TLP) loaded with siRNAs

Synthesis of TLPs and production of siRNA-TLPs was done exactly as recently described [44]. 10^5^ HeyA8 cells (infected with a luciferase lentivirus and a NucRed lentivirus (Essen Bioscience)) were injected i.p. into 6-week-old female NSG mice [44] following the Northwestern University Institutional Animal Care and Use Committee (IACUC)-approved protocol. The growth of tumor cells in the mice over time was monitored non-invasively using the IVIS^®^ Spectrum *in vivo* imaging system as recently described [44]. Each mouse of a treatment group was injected with 150 μl of either siNT-TLP or siCAG/CUG-TLP (1 μM stock).

### Data analyses

To determine the number of triplets, 6mer, 10mer or 19mer repeat sequences in the ORFs or 3’UTR of human or mouse genes, all ORF and 3’UTRs were extracted from the *Homo sapiens* (GRCh38.p7) or *Mus musculus* (GRCm38.p5) gene dataset of the Ensembl database using the Ensembl Biomart data mining tool. For each gene, only the longest deposited ORF or 3’UTR was considered. Custom perl scripts were used to identify whether each 3’UTR or ORF contained an identical match to a particular triplet, 6mer, 10mer or 19mer.

GSEA was performed using the GSEA v2.2.4 software from the Broad Institute (http://www.software.broadinstitute.org/gsea); 1000 permutations were used. Two list of 1846 survival and 418 nonsurvival genes were used as recently described [15, 45]. They were set as custom gene sets to determine enrichment of survival genes versus the nonsurvival control genes in downregulated genes from the RNA-seq data. Log(Fold Change) was used as the ranking metric. p-values below 0.05 were considered significantly enriched. The GO enrichment analysis shown was performed using all genes that after alignment and normalization were found to be at least 1.5 fold downregulated with an adjusted p values of <0.05, using the software available on http://www.Metascape.org and default running parameters. The data sets of HeyA8 cells with introduced siL3, shL1, shL3 or shR6 were recently described [15].

Sylamer analyses [25] were performed using the RNA-seq datasets from the HeyA8 cells transfected with siNT, siCAG/CUG or siCGA/UCG as recently described [15]. The analyses were performed using default settings. Enriched 6 or 10mer motifs were analyzed using either ORFs or the 3’UTRs sequences. Sylamer (version 12-342) was run with the Markov correction parameter set to 4. DAVID gene ontology analysis was performed using the tool at https://david.ncifcrf.gov/home.jsp and default settings.

### Statistical analyses

Two-way analysis of variances (ANOVA) were performed using the Stata 14 software to compare growth curves. One-tail student t-test was performed in the software package R to compare tumor load between treatment groups. Wilcoxon Rank Sum test was performed in R to compare IVIS signal between treatment groups. The effects of treatment on wild-type versus Drosha^-/-^ cells were statistically assessed by fitting regression models that included linear and quadratic terms for value over time, main effects for treatment and cell type, and two- and three-way interactions for treatment, cell-type and time. The three-way interaction on the polynomial terms with treatment and cell type was evaluated for statistical significance since this represents the difference in treatment effects over the course of the experiment for the varying cell types. All statistical analyses were conducted in Stata 14 (RRID:SCR_012763) or R 3.3.1 in Rstudio (RRID:SCR_000432) except for Pearson correlation analyses, which were performed using StatPlus 6.2.2.

## Acknowledgements

We are grateful to Dr. Evangelos Kiskinis for advice on trinucleotide repeat expansion diseases and Sarah Fazal for performing siRNA screens. We would like to thank Johannes Peter for help in creating the movies. The mouse *Ago1*-4 k.o. embryonic stem cells were a kind gift from Dr. Markus Hafner. This work was funded by training grants T32CA070085 (to M.P.) and T32CA009560 (to W.P. and K.M.M), and R35CA197450 (to M.E.P.). C.S.T. would like to thank the Department of Defense/Air Force Office of Scientific Research (FA95501310192) for grant funding, and grant funding from the National Institutes of Health/National Cancer Institute (R01CA167041). K.M.M acknowledges the Ryan Family, the Malkin Family, the Driskill Family, Chicago Baseball Charities Cancer Fellowship, The Northwestern University Feinberg School of Medicine Developmental Therapeutic Institute for financial and fellowship support.

## Author contributions

MEP and AEM designed and supervised the project; AEM, WP, QG, MP, CL, and BB performed research, and analyzed data; EB provided bioinformatics support and analyzed data; SC performed the siRNA screens; KMM and CST provided the siRNA-TLP particles; MEP wrote the manuscript, and all authors reviewed and approved the manuscript.

## Appendix files - Table of contents

**Appendix Figure S1.**
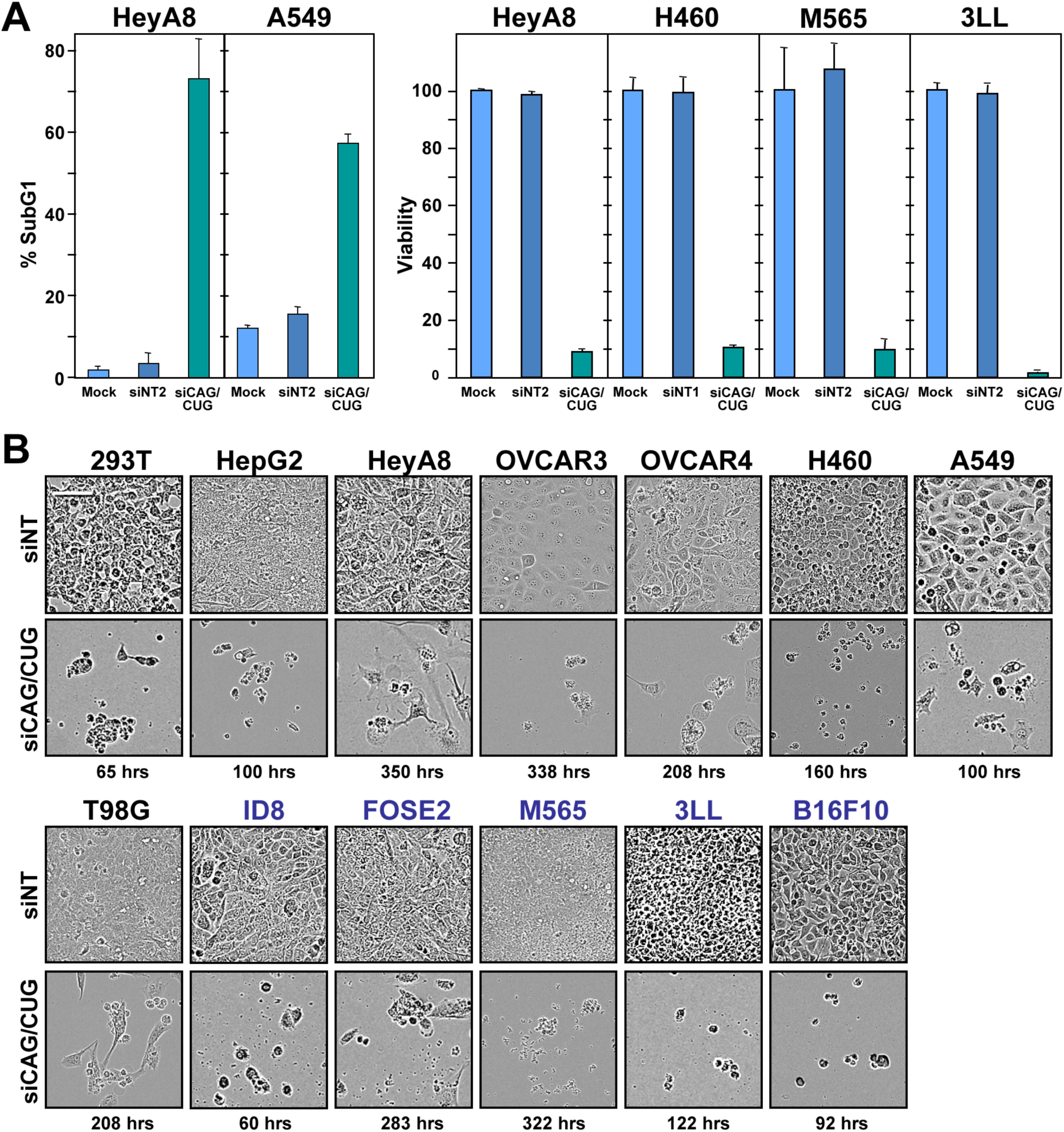
Morphological changes and cell death in cells transfected with siCAG/CUG. (A) *Left:* DNA fragmentation in HeyA8 and A549 cells 120/72 hours after transfection with indicated siRNAs. n = 3-4 technical replicates. *Right:* Viability of human and mouse cancer cell lines 96 hours after transfection with the indicated siRNAs. Values are mean −/+SD. n = 2 biological replicates, 3 technical replicates each. (B) Human (black text) and mouse (blue text) cell lines transfected with either siNT or siCAG/CUG. Time points after transfection of picture taking is given. Size bar = 100 μm.

**Appendix Figure S2.**
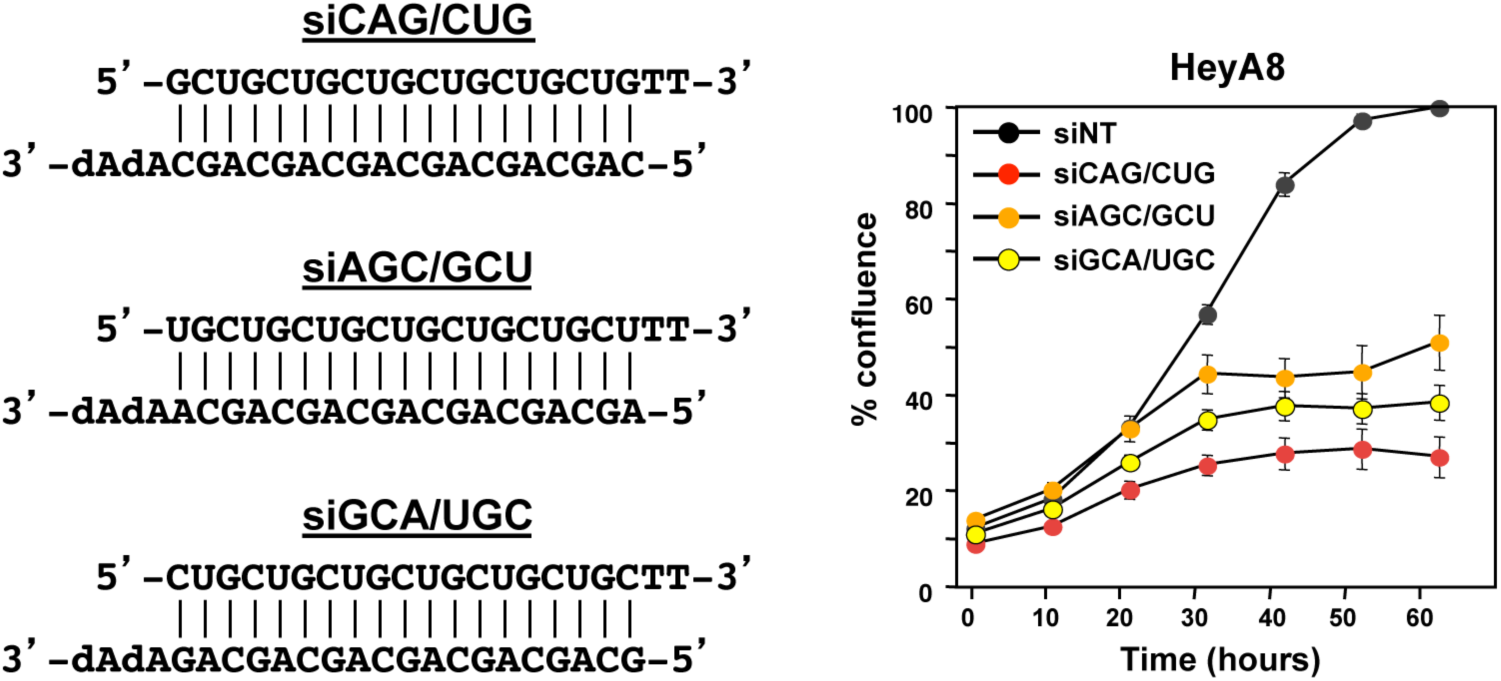
Properties of toxic TNR based siRNAs. *Left*: Sequences of the siCAG repeat in three different frames. *Right*: Confluence over time of HeyA8 cells transfected with 1 nM of either siNT or the three duplexes depicted on the left. Values are mean −/+SEM. n = 3 biological replicates, 4-8 technical replicates each.

**Appendix Figure S3.**
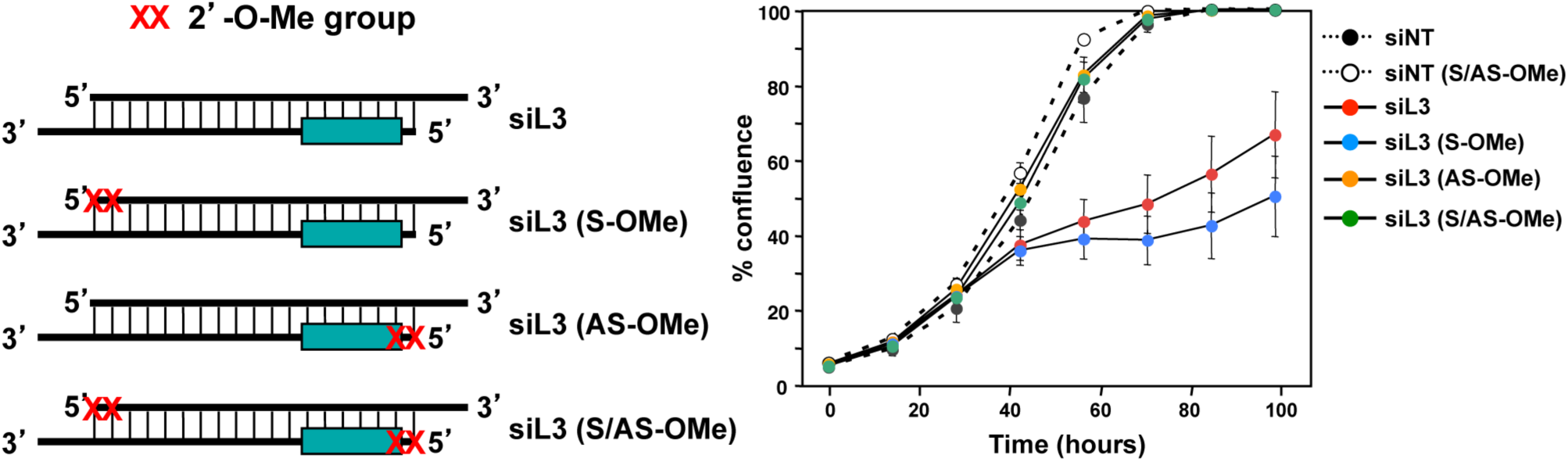
The toxicity of siL3 is solely based on its guide strand. *Left*: Scheme showing positions of the 2’O-methylation in either the passenger/sense (S) or the guide/antisense (AS) strand of siL3. The siL3 seed region is shown as a green box. *Right*: Confluence over time of HeyA8 cells transfected with 10 nM of the four duplexes depicted on the left and two similarly modified duplexes derived from siNT. Values are mean −/+SEM. n = 3 technical replicates.

**Appendix Figure S4.**
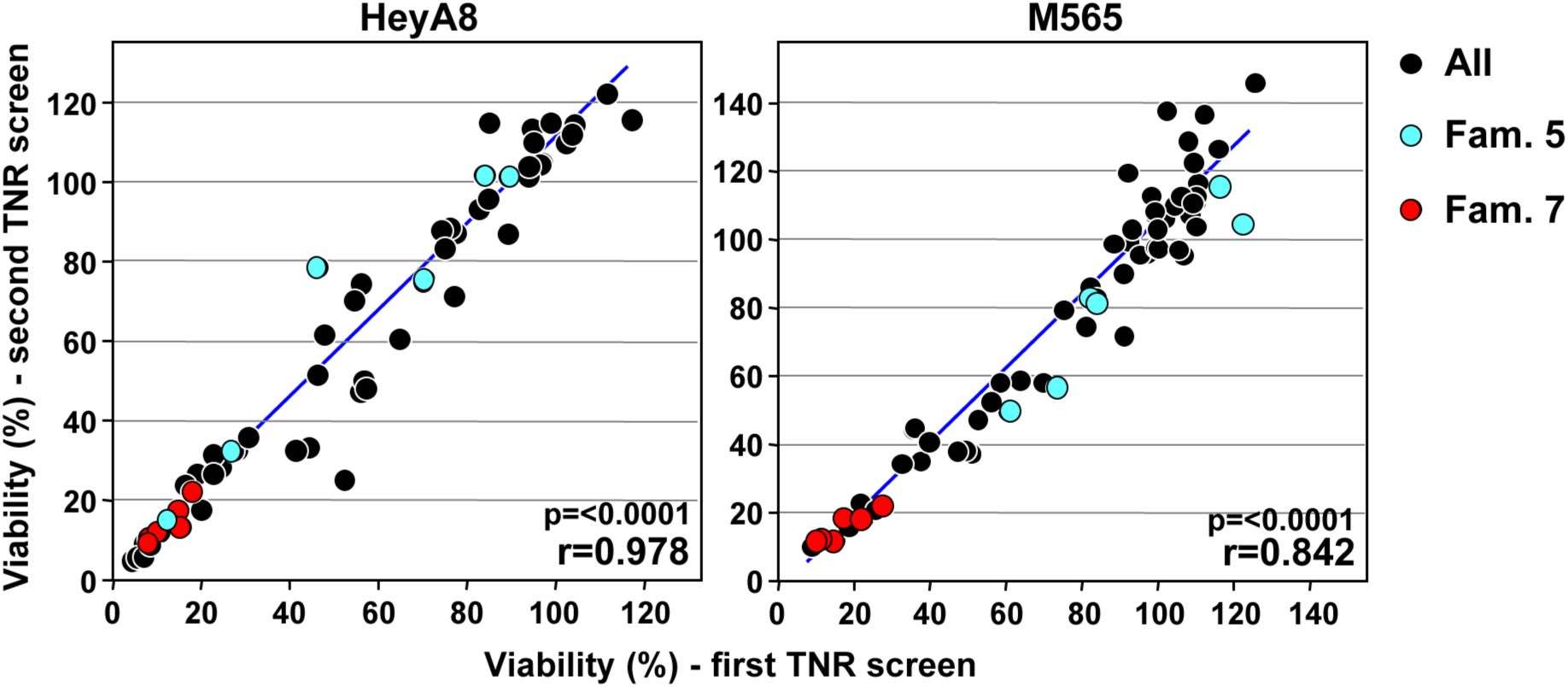
Reproducibility of the TNR siRNA screens. Variation between the two screens of the 60 TNR based siRNAs in HeyA8 (left) and in M565 (right) cells. The data points for the six members of TNR family 5 are labeled in blue those of family 7 are labeled in red. Pearson correlations and p-values are given.

**Appendix Figure S5.**
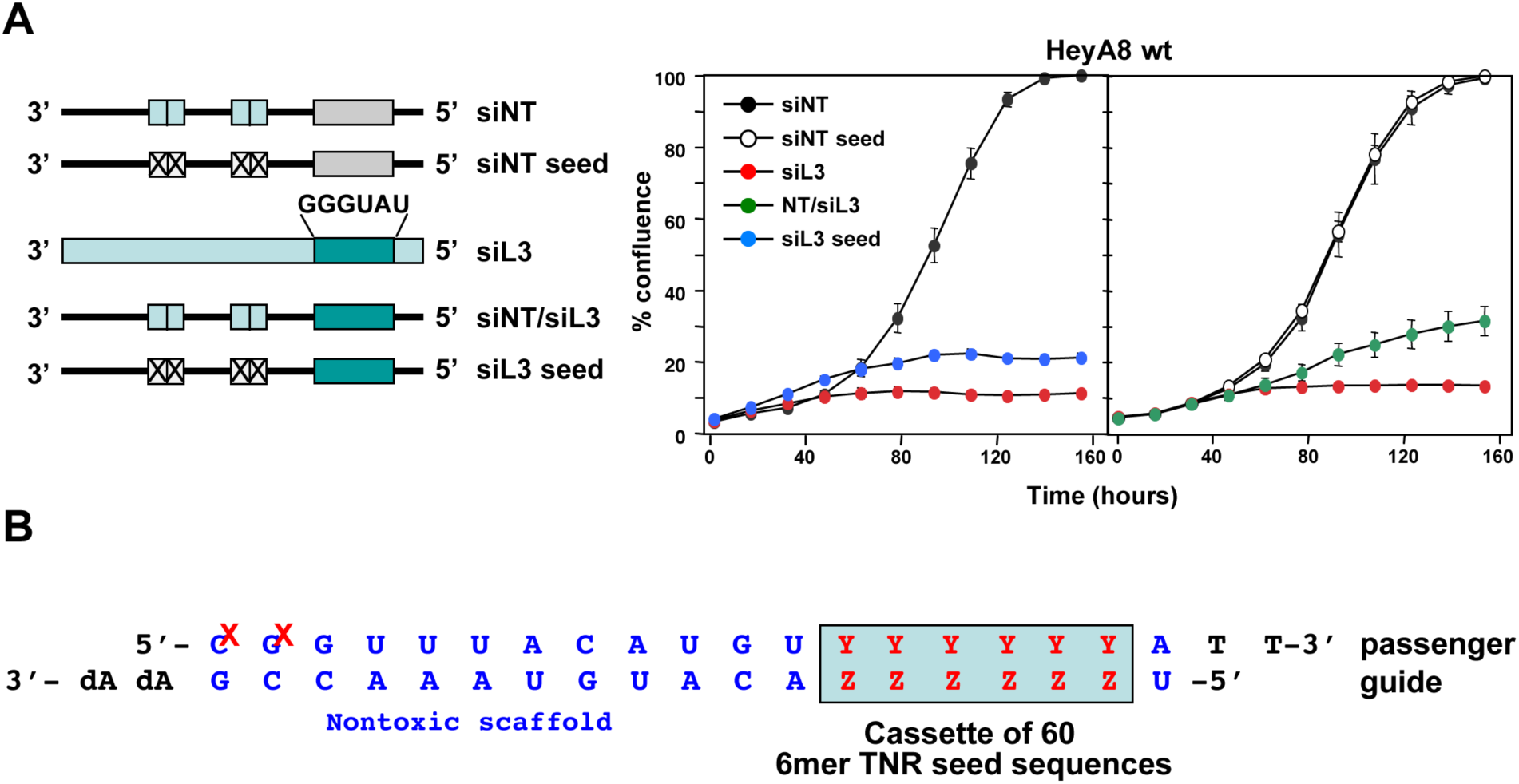
DISE inducing activity of siL3 is mostly based on its seed sequence. (A) *Left*: Scheme showing the different mutant siNT and siL3 duplexes used. The four light blue boxes indicate the four positions that in the nontargeting siRNA (siNT) were identical to the same positions in siL3. In siL3 the seed sequence is shown as a green box, and the siNT seed sequence is shown as a grey box. siNT/siL3 is a chimeric duplex comprised of the seed sequence of siL3 and the rest of siNT. In the siL3 seed duplex, the four siL3 positions in siNT were replaced with the complementary nucleotides (i.e. an G:C was changed to a C:G). *Right*: Confluence over time of HeyA8 cells transfected with 10 nM of the five duplexes depicted on the left. Values are mean −/+SEM. n = 6 technical replicates. (B) Schematic showing the design of the seed siRNAs tested in Fig 2D. Y and Z indicate the Watson-Crick complementary nucleotides of the 6mer seed. The two red Xs indicate the position of the 2’O-methylation in the passenger strand.

**Appendix Figure S6.**
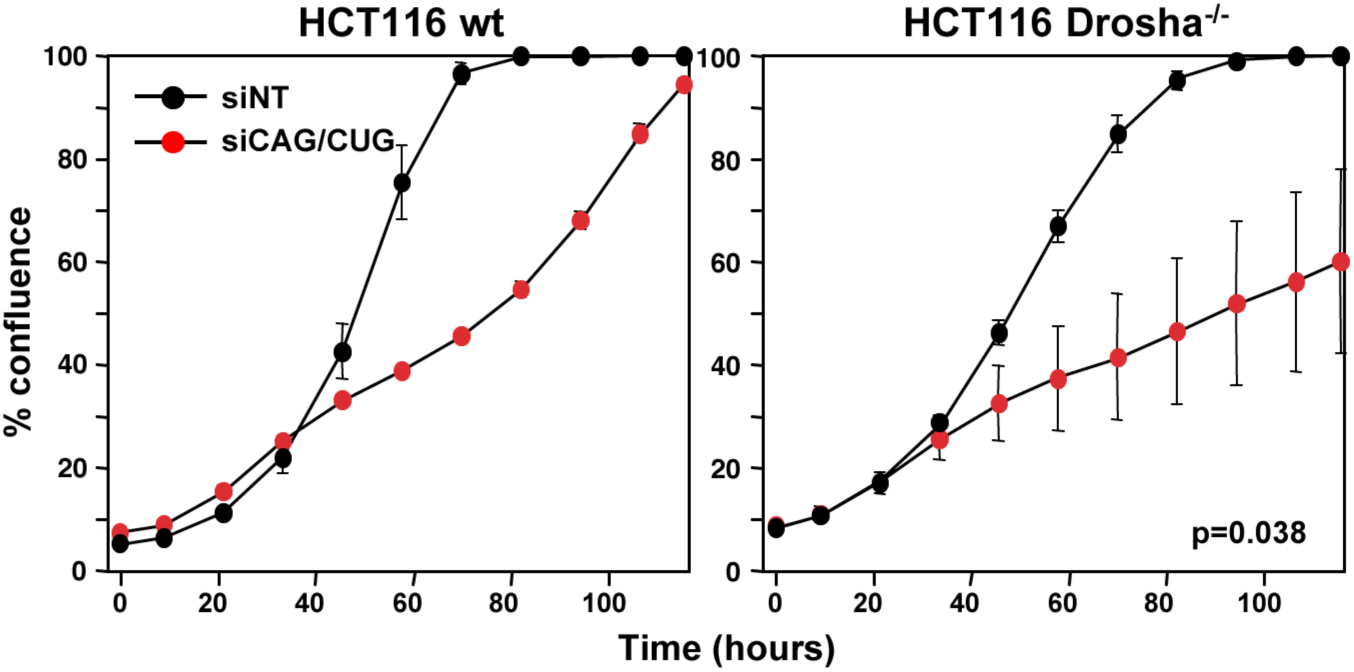
Increased sensitivity of HCT116 Drosha^-/-^ cells compared to HCT116 wild-type cells. Confluency over time of either HCT116 wt (left) or Drosha^-/-^ (right) cells transfected with 0.1 nM of either siNT or siCAG/CUG. Transfection efficiency of the two cell lines was similar as assessed by uptake of siGLO [1]. p-value according to polynomial distribution is given. Values are mean −/+SEM. n = 3 biological replicates, 3-4 technical replicates each.

**Appendix Figure S7.**
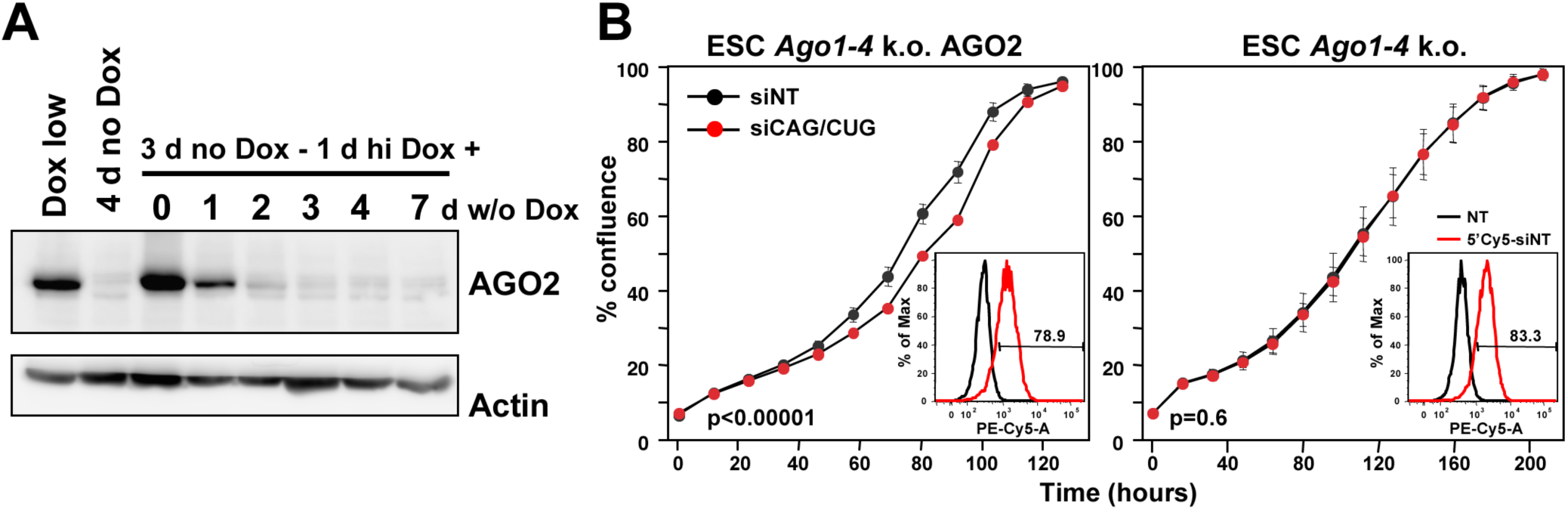
siCAG/CUG induced growth reduction of mouse embryonic stem cells requires *Ago2*. (A) Sensitivity of mouse embryonic stem cells to siCAG/CUG lacking expression of all four AGO proteins with the same cells in which we rendered the RISC functional by re-expression of human AGO2. Western blot analysis of *Ago1-4* k.o. mouse embryonic stem cells expressing a Tet inducible AGO2 protein. Lanes are as follows: 1: Low Dox; 2: 4 days without Dox; 3: 3 days without Dox - 1 day with high Dox; 4: 3 days without Dox - 1 day with high Dox - 1 day without Dox; 5: 3 days without Dox - 1 day with high Dox - 2 days without Dox; 6: 3 days without Dox - 1 day with high Dox - 3 days without Dox; 7: 3 days without Dox - 1 day with high Dox - 4 days without Dox; 8: 3 days without Dox - 1 day with high Dox - 7 days without Dox. Low Dox = 0.1 mg/ml; high Dox = 2.5 mg/ml. n = 2 biological replicates. (B) Confluence over time of *Ago1-4* k.o. cells with induced AGO2 with high Dox (left) or without Dox (right) after transfection with 5 nM siNT or siCAG/CUG. Two-way ANOVA is given. Equal transfection efficiency was established by transfecting cells with 5 nM of either siNT or 5’Cy5 labeled siNT followed by FACS analysis (inserts). Values are mean −/+SEM. n = 3 biological replicates, 4 technical replicates each. AGO2 expressing *Ago1-4* k.o. ESCs over-expressing human AGO2 showed a very low but reproducible susceptibility to siCAG/CUG. In contrast, *Ago1-4* k.o. cells were completely resistant to this form of toxicity, despite similar transfection efficiencies (see inserts) between *Ago1-4* k.o. cells and those expressing AGO2. The low sensitivity of the *Ago1-4* k.o./AGO2 cells could either be due to normal cells being less sensitive to this form of cell death or could point at a functional role of Ago family members other than Ago2 in this process.

**Appendix Figure S8.**
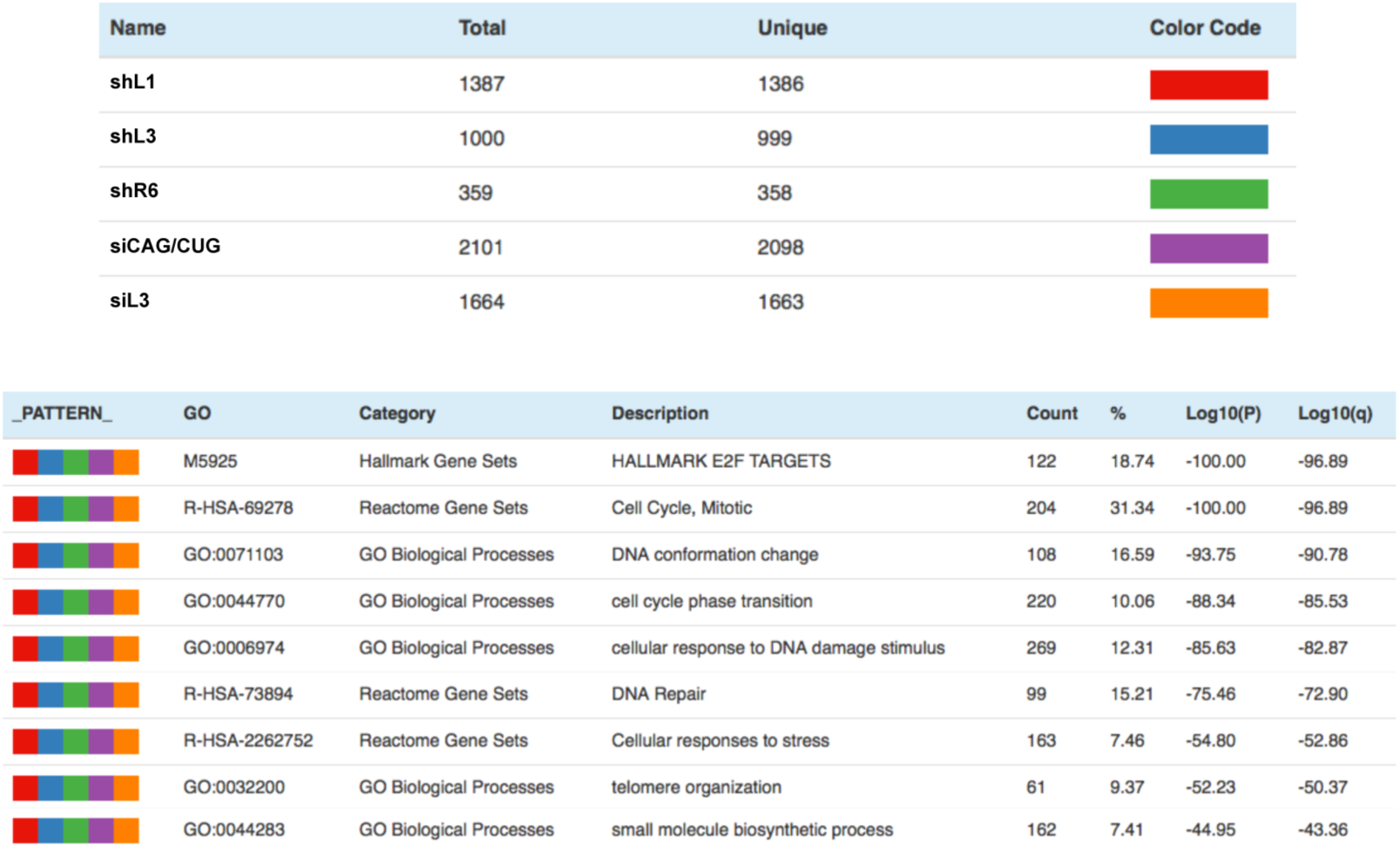
Similar pathways are affected in cells treated with siCAG/CUG and cells undergoing DISE. Metascape analysis of four RNA-Seq data sets of cells with introduced siL3, two CD95L derived shRNAs (shL1 and shL3), a CD95 derived shRNA (shR6), previously described [1], and the downregulated genes in cells transfected with siCAG/CUG. Shown are all GO terms enriched in all five data sets.

**Appendix Figure S9.**
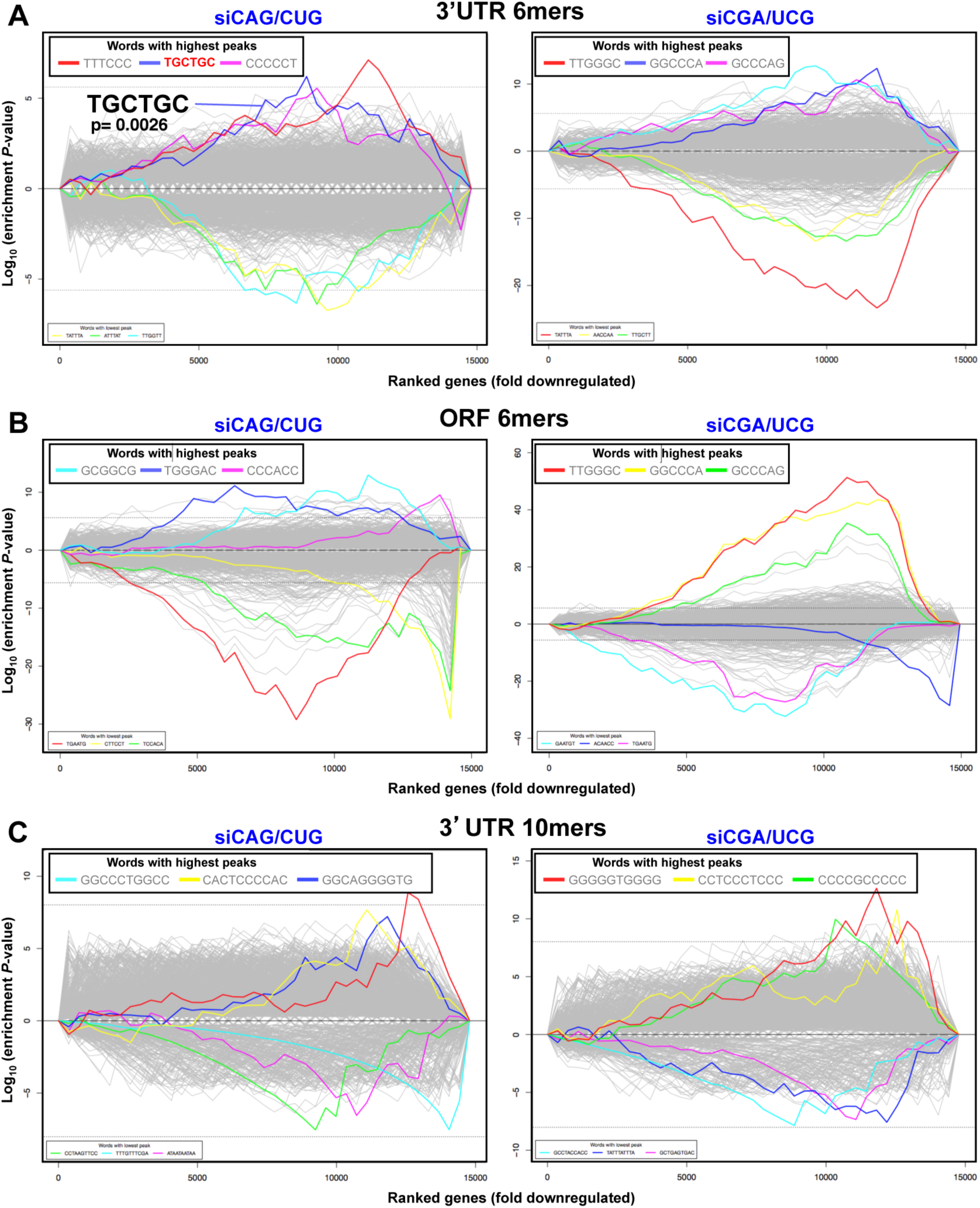
No significant enrichment of targeted seed sequences in the 3’UTR of downregulated genes. (A, B) 6-nucleotide Sylamer analysis of the 3’UTRs (A) or ORFs (B) of genes deregulated in HeyA8 cells transfected with either siCAG/CUG (left) or siCGA/UCG (right) ordered from most down-regulated to most up-regulated. The three most highly enriched sequences are shown. The 6- nucleotide motif in bold in the siCAG/CUG analysis corresponds to the reverse complement of position 2-7 in the 19mer siCAG strand. Bonferroni-adjusted p-value is shown. (C) 10-nucleotide Sylamer analysis of the 3’UTRs of genes deregulated in HeyA8 cells transfected with either siCAG/CUG (left) or siCGA/UCG (right) ordered from most down-regulated to most up-regulated. The three most highly enriched sequences are shown. Sequences without a p-value were either not significantly enriched or not at the top of the ranked list.

**Appendix Figure S10.**
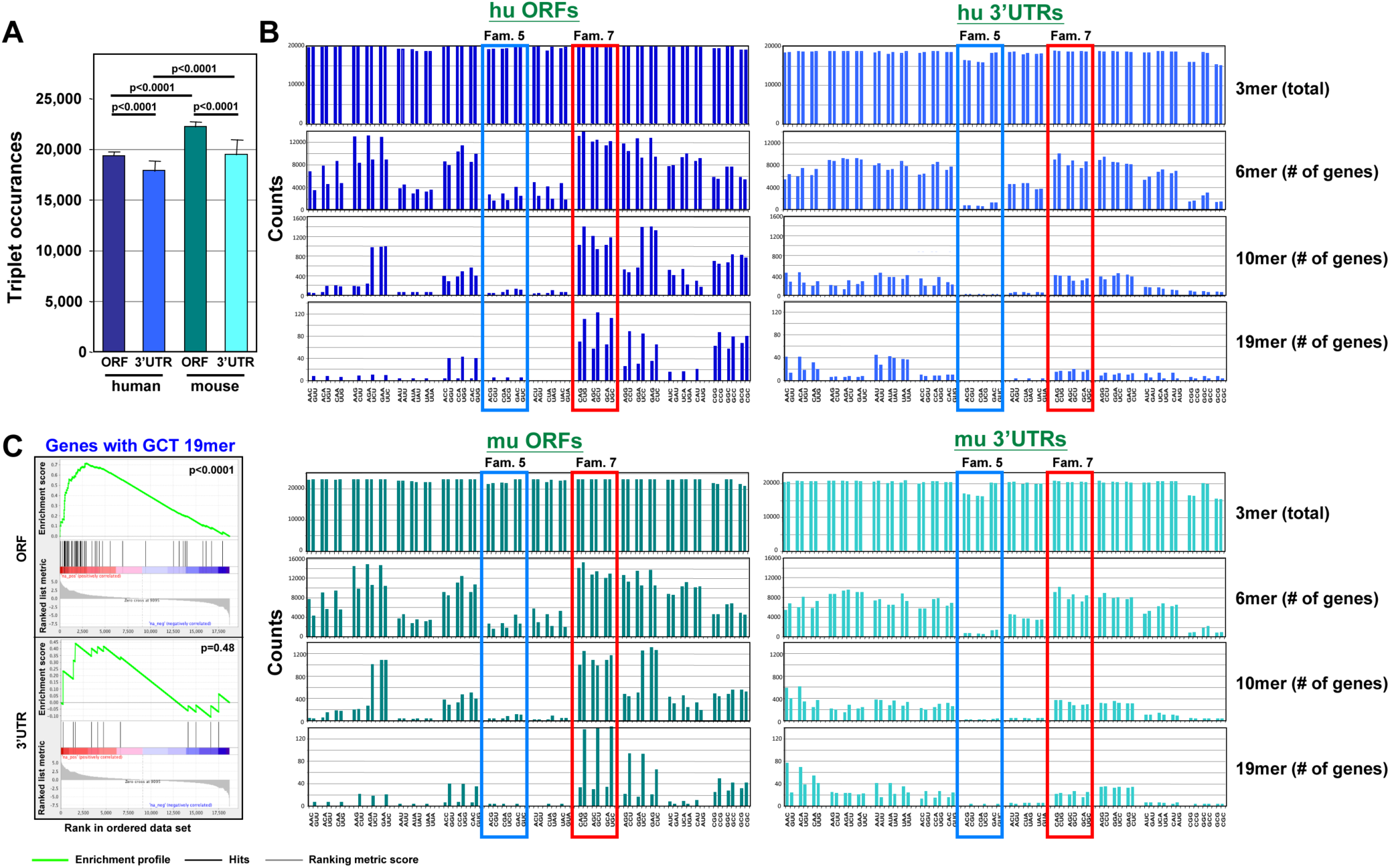
Vastly different frequencies of TNR expansions in the ORFs and 3’UTRs of human and mouse genomes. (A) Frequencies of all triplets combined in mouse and human ORFs and 3’UTRs. Values are mean −/+SD. Unpaired two tailed equal variance ttest was used to calculate p values. (B) *Top panels:* Frequencies of individual triplets in mouse and human ORFs and 3’UTRs. *Bottom three panels:* Number of genes containing 6mers, 10mers, or 19mers targeted by the 60 TNRs. Family 5 is boxed in blue and family 7 is boxed in red. (C) Gene set enrichment analysis for genes containing a GCTGCTGCTGCTGCTGCTG 19mer (targeted by the siCAG 19mer) in their ORFs (top) and 3’UTRs (bottom) in cells transfected with siCAG/CUG when compared to cells transfected with siNT. p values were calcuated using the Kolmogorov-Smirnov test.

**Appendix Figure S11.**
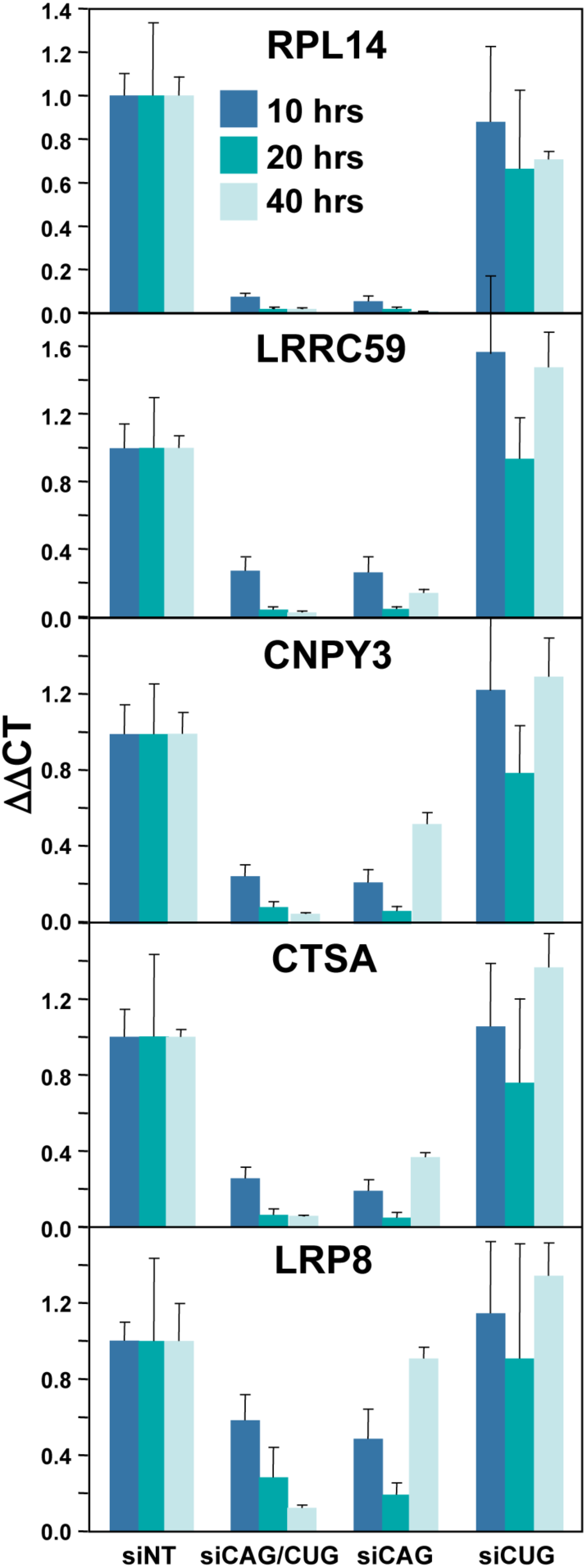
Efficient knockdown of CUG repeat containing genes in cells transfected with siCAG. HeyA8 cells were transfected with 1 nM of either siNT, siCAG/CUG, siCAG (with the CUG containing passenger strand modified by 2’O-methylation), or siCUG (with the CAG containing passenger strand modified by 2’O-methylation). RNA was quantified by real-time PCR. The genes are ranked according to their highest fold downregulation in the RNA Seq experiment. Values are mean −/+SD. n = 2 biological replicates (for siNT and siCAG/CUG), 3 technical replicates each.

**Appendix Figure S12.**
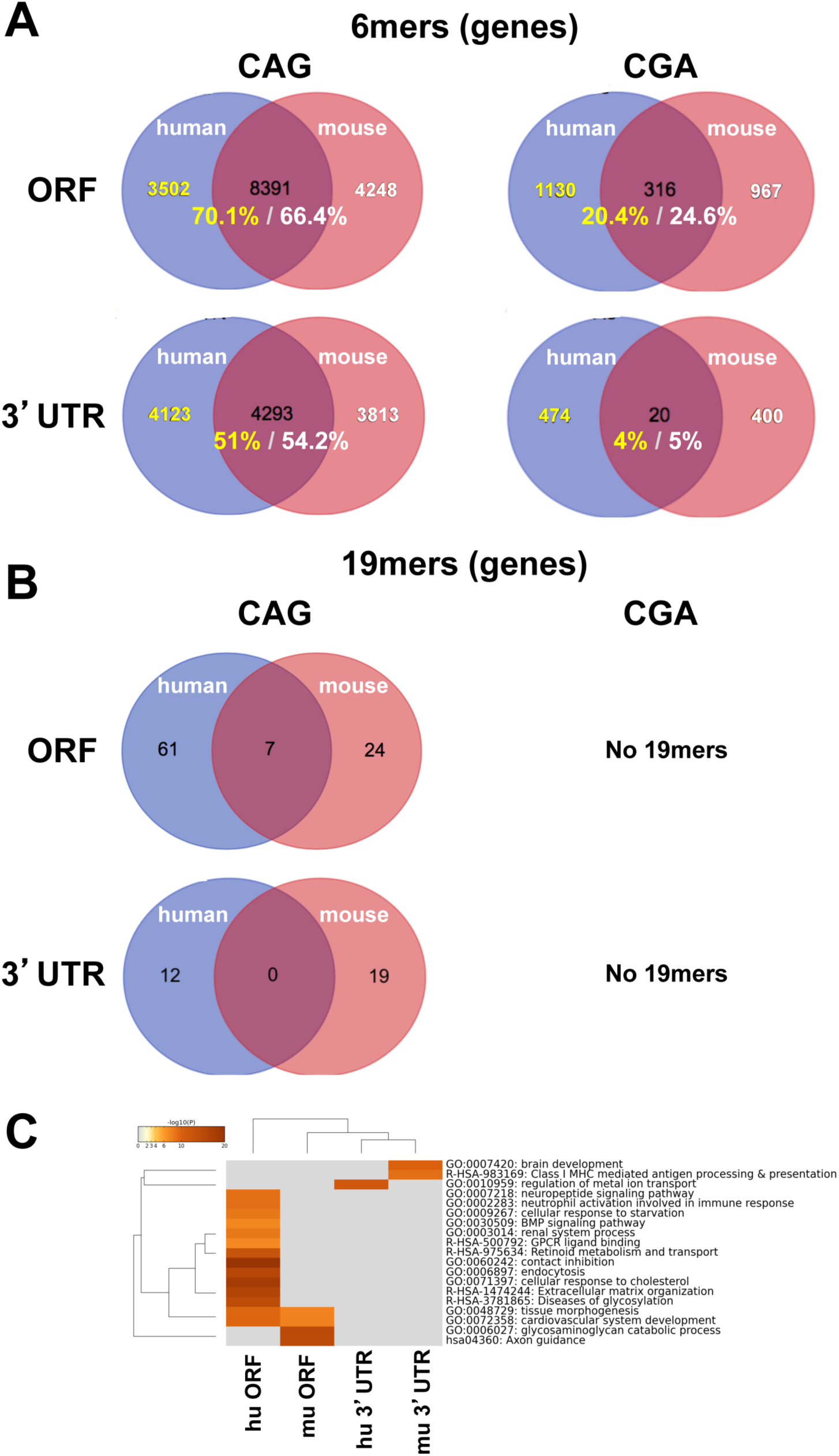
Genes containing the 19mer targeted by siCAG are poorly conserved between human and mouse. (A) Venn diagram of human and mouse ORFs and 3’UTRs containing the CAG (left) or CGA (right) nucleotide triplets. (B) Venn diagram of human and mouse ORFs and 3’UTRs containing the 19mer sequences completely complementary to the CAG (left) or CGA (right) based 19mer. (C) Metascape analysis of human and mouse ORFs and 3’UTRs containing the 19mer sequences completely complementary to the CAG 19mer.

**Appendix Figure S13.**
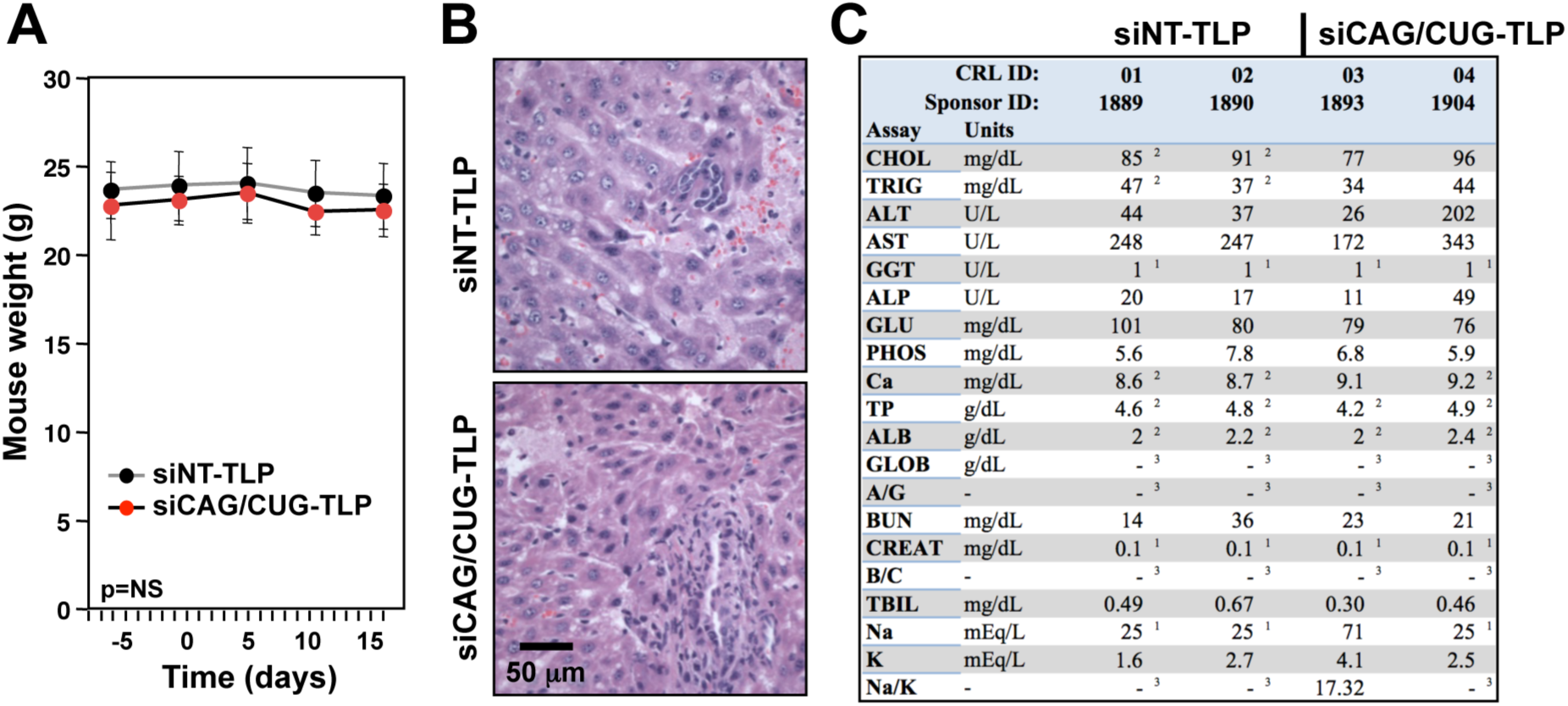
No adverse effects in mice treated with siCAG/CUG-TLPs. (A) Weight of the ten mice in treatment group 1 (see Fig 5D) over the course of the treatment. Values are mean −/+SD. NS, unpaired p-value not significant. (B) H&E stained liver sections of two of the mice that were treated with either siNT-TLP or siCAG/CUG-TLP on day 27 in the experiment shown in Fig 5E). (C) Serum analysis of the same two mice per treatment group. 1 = Sample assay value is less than the dynamic range. For most assays, the dynamic range low limit is reported. 2 = Sample was diluted for testing. Assay value for sample was below dynamic range, but results have been corrected for dilution. 3 = Assay is a calculated value. Either or both assay values used in the calculation were below the dynamic range of the assay, therefore no result is reported.

